# Donnan dominated ion homeostasis and the longevity of ischemic Na^+^-loaded dystrophic skeletal muscle

**DOI:** 10.1101/2020.11.20.391839

**Authors:** Catherine E Morris, Joshua J Wheeler, Béla Joos

## Abstract

The inherited muscle-wasting disease, Duchenne muscular dystrophy (DMD), renders skeletal muscle fibers (SMFs) Na^+^-overloaded, ischemic, membrane-damaged, cation-leaky, depolarized, and prone to myogenic firing. DMD fibers nevertheless survive up to 3 decades before succumbing to Ca^2+^-necrosis. The Ca^2+^-necrosis is explicable, the longevity is not. Modeling here shows that SMFs’ ion homeostasis strategy, a low-cost resilient Pump-Leak/Donnan feedback process we term “Donnan dominated”, underpins that longevity. Together, SMFs’ huge chloride-permeability and tiny sodium-permeability minimize excitability and pump costs, facilitating the outsized SMF pump-reserve that lets DMD fibers withstand deep ischemia and leaky channels. We illustrate how, as these impairments intensify, patients’ chronic Na^+^-overload (now non-invasively evident via Na^23^-MRI) would change. In simulations, prolonged excitation (→physiological Na^+^-overloading) and/or intense ischemia (→too little Na^+^-pumping) and accumulated bleb-damage (→too much Na^+^-leaking) eventually trigger Ca^2+^-overloading conditions. Our analysis implies an urgent need to identify SMFs’ pivotal small P_Na_, thereby opening new therapeutic remediation routes.

## INTRODUCTION

### Skeletal muscle fibers (SMFs) and dystrophin

SMFs are large long-lived multinucleate force-generating cells that are also subject to mechanical stresses. Their sarcolemmal bilayer, as for any plasma membrane, represents the fragile ~4 nm wide osmotic and electrical boundary across which electrical signals and ion homeostasis are mediated. In SMFs, bilayer tearing and bleb-damage (dissociation of bilayer from adherent cytoskeleton proteins) is minimized by dystrophin, an exceptionally long filamentous protein bound to actin, microtubules, lipids and more, and anchored at the massive transbilayer sarcolemmal dystroglycan complex (Campbell and Kahl 1989; Ibraghimov-Beskrovnaya et al 1992; Prins et al 2009; Khairallah et al 2012; Sarkis et al 2013; Belanto et al 2014; Dos Santos Morais et al 2018). Dystrophin acts as an entropic spring shock absorber (Le et al 2018), providing mechanical resilience while rendering the membrane skeleton “smart” through its interactions with various signaling proteins (Constantin 2014; Allen et al 2016). Sporadically, to compensate for accumulated wear and tear, adult SMFs regenerate; this is achieved by SMF fusion with stem (satellite) cells that have had a surge of dystrophin expression (Wang et al 2013; Dumont et al 2015; Chang et a 2016).

In the X-linked muscle-wasting disease, Duchenne muscular dystrophy (DMD), patients become non-ambulatory in their teens and live <3 decades dealing with wide-ranging ion homeostatic/metabolic impairment, most notably muscle ischemia (Allen et al 2016; Bishop et al 2018; Huong et al 2018; Gerhalter et al 2019). The chronic Na^+^-overload of dystrophic SM was first reported in 1957 (**Table 1**, item **1**). In 1988, Zubrzycka-Gaarn and colleagues reported that, unexpectedly, the DMD gene encodes not an “energy” enzyme but dystrophin, a membrane skeletal protein (Allen et al 2016), and dystrophin-minus mice, *mdx* (Ryder-Cook et al 1988), were introduced.

**TABLE 1.** Chronic Na^+^-overload of DMD patient muscles and *mdx* SMFs

| # | REFERENCE | [Na<sup>+</sup>]<sub>i</sub> | NOTES |
| --- | --- | --- | --- |
| 1 | 1955 – Horvath et al | [Na<sup>+</sup>]<sub>i</sub> higher (& [K<sup>+</sup>]<sub>i</sub> lower ) in dystrophic than in normal muscle fibers | (Horvath et al 1955 is referenced in Rudman et al 1972); biopsy, elemental analysis; fibers of 20 **patients** with unspecified muscular dystrophies |
| 2 | 1993 – Dunn et al | diaphragm, gastrocnemius<br>control: 13.0, 13<br>**mdx**: 23.5, 24 | Two techniques: Na<sup>+</sup>-electrode for diaphragm, cyto-volumetrics plus serum and bulk muscle Na values for gastrocnemius. Mice. |
| 3 | 2008 – Hirn et al | 1.4X more <sup>22</sup>Na<sup>+</sup> uptake in *mdx* than in control | <sup>22</sup>Na<sup>+</sup> uptake. Tetrodotoxin reduces uptake in both and makes them identical (see their Fig 1). Mice. |
| 4 | 2011 – Miles et al | control: 11.5 mM<br>**mdx**: 22.5 mM | Na<sup>+</sup>-dye measurements. Mice. |
| 5 | 2011, 2012 – Weber et al | <u>Total sodium content</u>:<br>volunteers: 25-26 mM<br>DMD **patients**: 38 mM<br><u>Normalized <sup>23</sup>Na IR (*)</u><br>volunteers: 0.50<br>DMD **patients**: 0.75 | <sup>23</sup>Na-MRI. German **patient** cohort.<br>Hypothesis that DMD edema is an osmotic effect due to ↑ myoplasmic Na<sup>+</sup>, **not inflammatory** extracellular edema as generally thought.<br>(* IR or “inversion recovery”; a means of assessing intracellular Na<sup>+</sup> levels) |
| 6 | 2012 – Lehmann-Horn et al | sustained small decrease of cytoplasmic Na<sup>+</sup> (& H<sub>2</sub>O) overload (n=1) | <sup>23</sup>Na-MRI - pilot study - prolonged “off-label” eplerenone treatment, one 22-yr-old female DMD **patient** |
| 7 | 2014 – Altamirano et al | Vastus lateralis<br>control: 8 mM<br>**mdx**: 18 mM | Na<sup>+</sup>-electrodes. Shear-stress stimulation of NO-pathway reduced *mdx* [Na<sup>+</sup>]<sub>i</sub> to 10 mM without altering control level (see their Fig 1B). Mice. |
| 8 | 2014 – Burr et al | control: 5.3 mM<br>**mdx**: 7.3 mM | Na<sup>+</sup>-electrodes (see their Fig 10). Mice. |
| 9 | 2017 – Glemser et al | ~20% drop in Na<sup>+</sup> & ~15% drop in muscle H<sub>2</sub>O overloads (n=1) | <sup>23</sup>Na-MRI - pilot study - 6 months “off-label” eplerenone treatment, one 7-yr-old male DMD **patient** (see their Table 1) |
| 10 | 2019 – Gerhalter et al | <u>Total sodium content</u>:<br>volunteers: 16.5 mM<br>DMD **patients**: 26 mM<br><u>Normalized <sup>23</sup>Na IR</u><br>volunteers: 0.69<br>DMD **patients**: 0.47 | <sup>23</sup>Na-MRI - DMD **patient** cohort in France - authors emphasize that Na<sup>+</sup>-loading is regularly observed in patients with DMD even in absence of water T<sub>2</sub> increases (see also Gerhalter et al 2017) |

Mechanistic clarity on how DMD’s lack of dystrophin engenders persistent skeletal muscle ischemia is still a work-in-progress (e.g., Timpani et al 2015; Moore et al 2020) but the ischemia has multiple facets (**Figure 1A,B,C**). Early *mdx* SMF membrane studies showed that expression of Na^+^/K^+^-ATPase is not impaired, with both Anderson (1991) and Dunn et al (1995) reporting pump-protein at above-normal levels. Anderson’s work did, however, include findings consistent with “pump-strength” sub-normality, the supra-normal pump-protein-density notwithstanding; *mdx* sarcolemma has abnormal phospholipid profiles, consistent with the cumulative *mdx* bleb-damage from mechanical strain, reactive oxygen species and ↑Ca^2+^ (Petrof et al 1993; Dudley et al 2016) and consistent with pump dysregulation (see Hossain and Clarke 2019).

**Figure 1.**
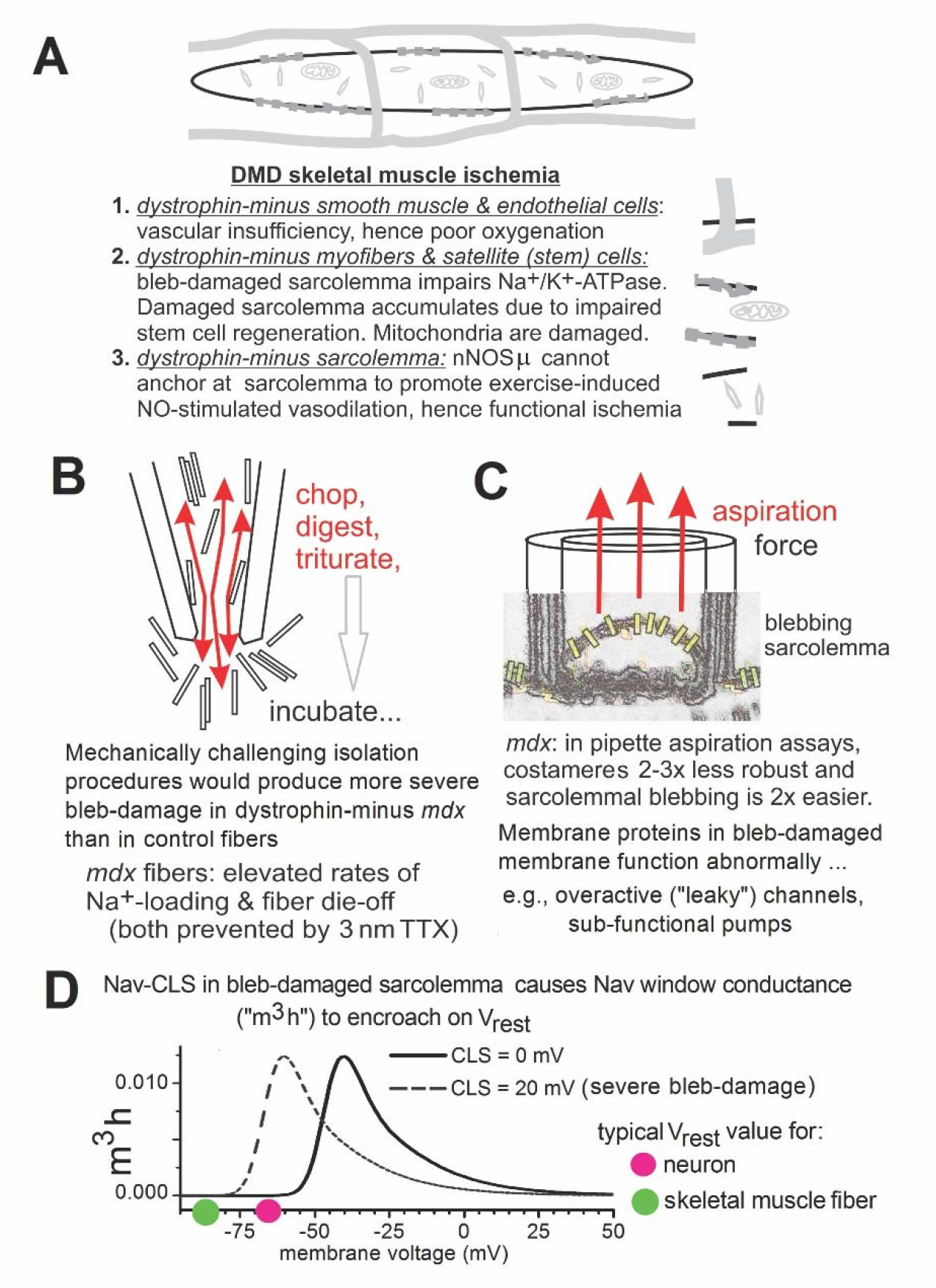
Dystrophic muscle fibers, ischemia and blebdamaged sarcolemma. **A.** SMF ischemia in DMD has several distinct causes **1.** Vascular insufficiency (Verma et al 2019; Podkalicka et al 2019). **2.** Malfunctioning pumps in bleb-damaged sarcolemma that accumulates because DMD fibers’ regenerative capabilities are impaired (Chang et al 2016; Shattock & Matsuura 1993; Williams and Bloch, 1999; Whitehead et al 2010; Juel et al 2015; Lansman 2015; Burr and Molkentin 2015; McElhanon & Bhattacharya 2018; Murphy et al 2019). **3.** Exertion-induced functional ischemia (Sander et al 2000; Allen et al 2016). **B.** Excessive fiber-isolation-induced bleb-damage to fragile (dystrophin-minus) Nav-bearing sarcolemma of *mdx* fibers would explain their “leaky” Nav1.4 channels (Hirn et al 2008; see Morris, 2018). **C.** Depiction of García-Pelagio et al (2011) pipette aspiration tests that quantified *mdx* fiber susceptibility to bleb-damage; bleb-damage signifies loss of non-covalent binding (adhesion) between bilayer and membrane skeleton proteins (e.g., F-actin) that fosters, e.g., decay of cell-mediated structural features of the bilayer (Methfessel et al 1986; Sheetz et al 2006; Phillips et al 2009; Lundbaek et al 2004, 2010; Morris 2012; Ameziane-Le Hir et al 2014; Cornelius et al 2015; Petrov et al 2017; Burden et al 2018). That in turn alters conformational stabilities of membrane proteins (Morris and Horn 1991; Morris et al 2012A,B; Zhang et al 2000; Zhang and Hamill 2000; Burr and Molkentin, 2015). Rectangles signify ion transporting proteins that malfunction in damaged sarcolemma (e.g., Na^+^/K^+^-pumps, Na^+^-permeant channels). **D**. For the computational Nav channel used here, window conductance (m^3^h(Vm)) before and after a 20 mV Nav-CLS or Nav Coupled-Left-Shift. Explanation: SMF-Nav and neuronal-Nav channels undergo irreversible hyperpolarizing shifts in bleb-damaged membrane (Boucher et al 2012; Morris and Joos 2016; Joos et al 2016). In Hodgkin-Huxley (H-H) terms, fast activation (m^3^) and inactivation (h) both shift “leftward” by the same amount (“CLS”).

During muscle exertion, dystrophin-dependent signaling normally ensures adequate microcirculation as follows: nNOSμ, a nitric oxide(NO)-synthetase bound to the dystrophin-dystroglycan complex, responds to exertion by locally increasing vasodilation-inducing NO (Sander et al 2000; Allen et al 2016). Lacking dystrophin to position nNOSμ, DMD fibers experience “functional ischemia” (Asai et al 2007; Thomas 2013; Altamirano et al 2014; Nelson et al 2015; Rebolledo et al 2016; Percival 2018) so contractile activity dangerously exacerbates DMD fibers’ persistent ischemia. Though clinically, “ischemia” means inadequate blood-flow, for modeling here it will signify sub-normal (<100%) Na^+^/K^+^-ATPase ***<u>strength</u>*** from any cause.

The DMD/*mdx* sarcolemma is mechanically vulnerable not only to bleb-damage, but to bilayer micro-tears and rupture (**Figure 1B,C**) (Petrof et al 1993, 2002; Menke and Jockusch 1991,1995; Dudley et al 2006; García-Pelagio et al 2011; Allen et al 2016; Houang et al 2018). Overexpression of the dystrophin-related protein, utrophin, substitutes only partially (e.g. Belanto et al 2014). Though microtears occur easily in DMD/*mdx,* fast exocytosis-based repair plus endocytic retrieval of resealed-patches functions normally (McNeil and Steinhardt 2003; Hody et al 2019; Barthélémy et al 2018; Corrotte et al 2013; Andrews et al 2014). Though myogenic contractures are extra-perilous for *mdx* fibers (Claflin and Brooks 2008) and though fiber-stretch injury can leave *mdx* fibers depolarized and inexcitable, recovery occurs within days or weeks (Anderson 1991; Call et al 2013; Pratt et al 2015; Baumann et al 2020). The fragility of DMD/*mdx* fibers is manifest here, along with the remarkable resilience of SMF ion homeostasis.

### Ion homeostasis and DMD SMFs

Of the many muscular dystrophies, DMD is not only the commonest, but the most severe (Datta and Ghosh 2020). Nevertheless, DMD fibers can survive 20-30 years, progressively more debilitated by chronic ischemia, plagued by sarcolemmal damage. Though sarcolemmal tear-repair is a must, defective-repair myopathies (dysferlin muscular dystrophies) are less severe than DMD (Bansal et al 2003; Bittel et al 2020). What pre-disposes even DMD-stricken SMFs for longevity, we posit, is their evolutionarily-ancient energy-efficient ion homeostatic strategy.

As **Figure 8** and its legend summarize, a companion paper (Joos & Morris submitted) explains the distinctive Donnan dominated strategy adopted by SMFs, contrasting it with the better-known (Dijkstra et al 2016) Pump-leak dominated strategy of neurons. SMFs are low duty-cycle excitable cells (i.e., mostly at V_rest_) whose inexpensive steady-state is achieved by exploiting impermeant myoplasmic anions (Donnan effectors) in passive collaboration with [big P_Cl_] (ClC-1-based). This stabilizes a V_rest_ that, thanks to exceedingly [small I_Naleak_], is deeply hyperpolarized. The [small I_Naleak_] simultaneously fosters large pump-reserves that ensure resumption of steady-state even after abnormally large perturbations.

Here our aim is: **1)** to detail, with reference to the dystrophin-minus deficits of DMD-SMFs (ischemia and sarcolemmal damage), how their Donnan dominated P-L/D strategy safeguards such fibers, and **2)** to explore how leaky Na^+^-permeant channels (in damaged sarcolemma) would contribute to the chronic Na^+^-overload, myogenic contractures and eventual susceptibility to Ca^2+^-necrosis that besets DMD fibers.

When pump-strength falls so low that the P-L/D feedbacks fail, Donnan forces become lethal through irreversible osmotic (“cytotoxic”) swelling. This is how neurons die in cortical spreading depression (Rungta et al 2015; Dijkstra et al 2016; Dreier et al 2018) but for DMD fibers, Ca^2+^-necrotic processes typically intervene (Burr & Molkentin 2015; Timpani et al 2015; Murphy et al 2019). In DMD fibers, cause-of-death assignment is complicated by ischemia impacting both Ca^2+^ (Smith et al 2013) and ion (Na^+^/K^+^) homeostasis. With SMF ion homeostasis not widely understood even for healthy fibers, attention has mostly focused on proximate causes of DMD fiber-loss monitored via Ca^2+^-dyes, as when fiber-stretch or repeated myogenic action potentials (APs) trigger catastrophic Ca^2+^ influx and contractures (Yeung et al 2005; Claflin & Brooks 2008). Therapeutically it would be preferable, however, to intervene, where possible, upstream of lethal Ca^2+^-dysregulation.

Like healthy SMFs, DMD fibers rely on E_Na_ for APs and to fuel Na^+^-transporters (glucose, pH, multiple ions, Ca^2+^, etc (Usher-Smith et al 2009)). The chronic Na^+^-overload of DMD fibers (**Table 1**) is, by definition, non-lethal (Weber et al 2011,2012). That being said, when reverse operation of Na^+^/Ca^2+^-exchangers causes *mdx* fiber death, diminished E_Na_ is a pre-requisite (Burr et al 2014). Here again, what triggers Ca^2+^-necrosis is severe upstream Na^+^-overload. Though Ca^2+^-overload typically proves lethal sooner than cytotoxic swelling, fixing Na^+^ homeostasis could be most fruitful therapeutically.

Reviewing the consensus view (Allen et al 2016) that Ca^2+^-necrosis underlies DMD fiber loss, Burr and Molkentin (2015) write that “given the known mechanical defects within the dystrophic plasma membrane...alterations in calcium and sodium levels likely stem...from excessive activation of various channels and exchangers”. Implicit, by elimination, is the assumption that “too much Na^+^-leak” across bleb-damaged sarcolemma explains DMD Na^+^-overload. Our P-L/D analysis indicates, however, that “too little Na^+^-pumping” is also critical for DMD Na^+^-overload.

### Addressing the basis of chronic Na^+^-overload via P-L/D modeling

For almost 70 years ion homeostatic dysfunction has been recognized as a DMD trait, but this has yet to be addressed through a theoretical framework. Chronic Na^+^-overload was first reported for dystrophic muscle in the 1950s and is presently monitored non-invasively in patients via ^23^Na-MRI (**Table 1,** items **5**,**9**, **10**). Ion homeostatic impairment of Vm regulation presents clinically as myogenic contractility (Ishpekova et al 1999; Nojszewska et al 2017). Muscle tissue edema (versus SMF edema) in DMD has been confusing but concurrent ^23^Na-MRI and proton-MRI (to gauge fiber-H_2_O) (Weber et al 2011; Gerhalter et al 2017,2019) is providing greater clarity. Quantifying muscle ischemia has been problematic, but that is changing (Zhang et al 2020; Deitz et al 2020)).

Our theoretical framework is the ultra-simple P-L/D excitable SMF model, SM-CD (Skeletal Muscle Charge Difference; Joos & Morris submitted), with specified modifications to mimic the DMD condition. Two classes of “leaky” Na^+^-permeant channels expected in damaged sarcolemma are tested and DMD chronic ischemia is ascribed, partly, to non-functional pump-protein in damaged sarcolemma (**Figure 1A**). We show both how ischemic membrane-damaged DMD fibers could remain viable, and how their typical demise (via Ca^2+^-necrosis) relates to treacherous P-L/D thresholds. In compartment syndrome, similar issues arise, albeit far more rapidly than for DMD. Commonalities include profound ischemia, muscle damage and the imperative to avoid irreversible thresholds (Johnstone & Ball 2019; Tatman et al 2020). Dooley and Chiasson (2014; see too, Siegel 1992) argue, interestingly, that DMD patients’ sustained limb contractures can secondarily produce (fasciotomy-relieved) compartment syndrome.

Cell and molecular therapies are emerging for DMD (e.g. Datta et al 2020), concurrent with improved non-invasive monitoring of patients’ ion homeostatic indicators (Mankodi et al 2017; Bishop et al 2018; Gerhalter et al 2019; Ropars et al 2020; Meng et al 2020; Pennati et al 2020). Whether Na^+^-overload in DMD patients indicates fiber swelling that “is likely osmotically relevant” and thus “may contribute to the progressive muscle degeneration” (Weber et al 2011,2012) has been unclear. While recent patient measurements confirm the Na^+^-overload (Gerhalter et al 2017, 2019) they find it to be mostly non-osmotic. Our P-L/D-based analysis predicts this. As ischemia deepens, it shows, SMF Na^+^-overload would remain essentially non-osmotic till the attendant ischemic depolarization triggered myogenic firing. P-L/D analysis explains both tolerance of anoxic episodes and the limits to that tolerance, showing how over-stimulation plus membrane damage become fiber-lethal.

Our findings constitute a low-resolution conceptual map for the major contours of DMD’s ion homeostatic impairment. In future, SM-CD could be augmented with Na^+^-transporters, Nav1.4’s slow-gating kinetics, pump char acteristics specific to SMFs, and more, to enhance its sophistication for biomedical research. **<u>To recap</u>: 1)** DMD fibers are chronically ischemic, Na^+^-loaded, leaky to Na^+^ and overly-contractile. **2)** DMD fibers can survive for decades**. 3)** Non-invasive monitoring of patients’ ion homeostatic indicators is improving sharply just as potential therapies are coming on-line. **4)** SMFs use a [big P_Cl_][small I_Naleak_] or Donnan dominated strategy for ion homeostasis and an accessible theoretical framework is now available. **5)** We show how robust ion homeostasis safeguards SMFs during deepening ischemia and/or worsening Na^+^-leaks; their [big P_Cl_][small I_Naleak_] strategy, we argue, substantially explains DMD fibers’ remarkable longevity.

## RESULTS

### Overview

SM-CD, a [big P_Cl_][small I_Naleak_] ion homeostasis (P-L/D) model for excitable SMFs is used to address the issue of DMD longevity. SMF’s [big P_Cl_] is mostly its well-studied (SMF-specific) channel, ClC-1 (Pedersen et al 2016; Park and Mackinnon 2018; Wang et al 2019). Curiously, though ClC-1’s role in passively stabilizing SMF’s hyperpolarized V_rest_ is well understood, its role in SMF ion homeostasis *per se* goes unrecognized (Jentsch and Pusch 2018; Jeng et al 2020). A recurring theme in the Results is the pivotal contribution of [small I_Naleak_]. The small P_Na_ through which [small I_Naleak_] flows is unidentified.

Since DMD fibers are chronically ischemic, we first itemize benefits that ischemic SM-CD gleans from the P_Na_ and P_Cl_ components of “[big P_Cl_][small I_Naleak_]”. The full excitable SM-CD ensemble --P_Na_, P_K_, P_Cl_, pump, H_2_O, Donnan effectors, Nav, Kv, C_m_--is then examined in contexts relevant to DMD (bouts of anoxia/recovery, ischemia at varying intensity, damage-induced Na^+^-leaks through cation and Nav channels, and over-stimulation). Globally this reveals: extreme insult/injury is needed to overwhelm the Donnan dominated P-L/D system. Demise occurs when, Na^+^-influx finally exceeds the pump’s capacity (pump-reserve), eliciting myogenic APs and cytotoxic swelling, i.e., *in silico* death by [Na^+^+Cl^-^+H_2_O]-overload. *In situ,* a Ca^2+^-necrosis *coup de grâce* would likely supersede irreversible swelling. Because ^23^Na-proton-MRI is being used to monitor DMD patients’ ion homeostatic indicators, the last figure shows that myoplasmic Na^+^-overloads deemed “safe” 1) are “non-osmotic” but 2) vary substantially with membrane-damage and ischemic intensity.

### Syncytial SMFs are even more efficient than depicted by SM-CD

For physiological perspective, comparisons are made in some instances between ion fluxes across the 2000 μm^2^ of SM-CD and CN-CD (CN-CD=Cortical Neuron Charge Density; initial Vol_cell_ and pump strengths are also identical, see **Table 2**).

**Table 2.**
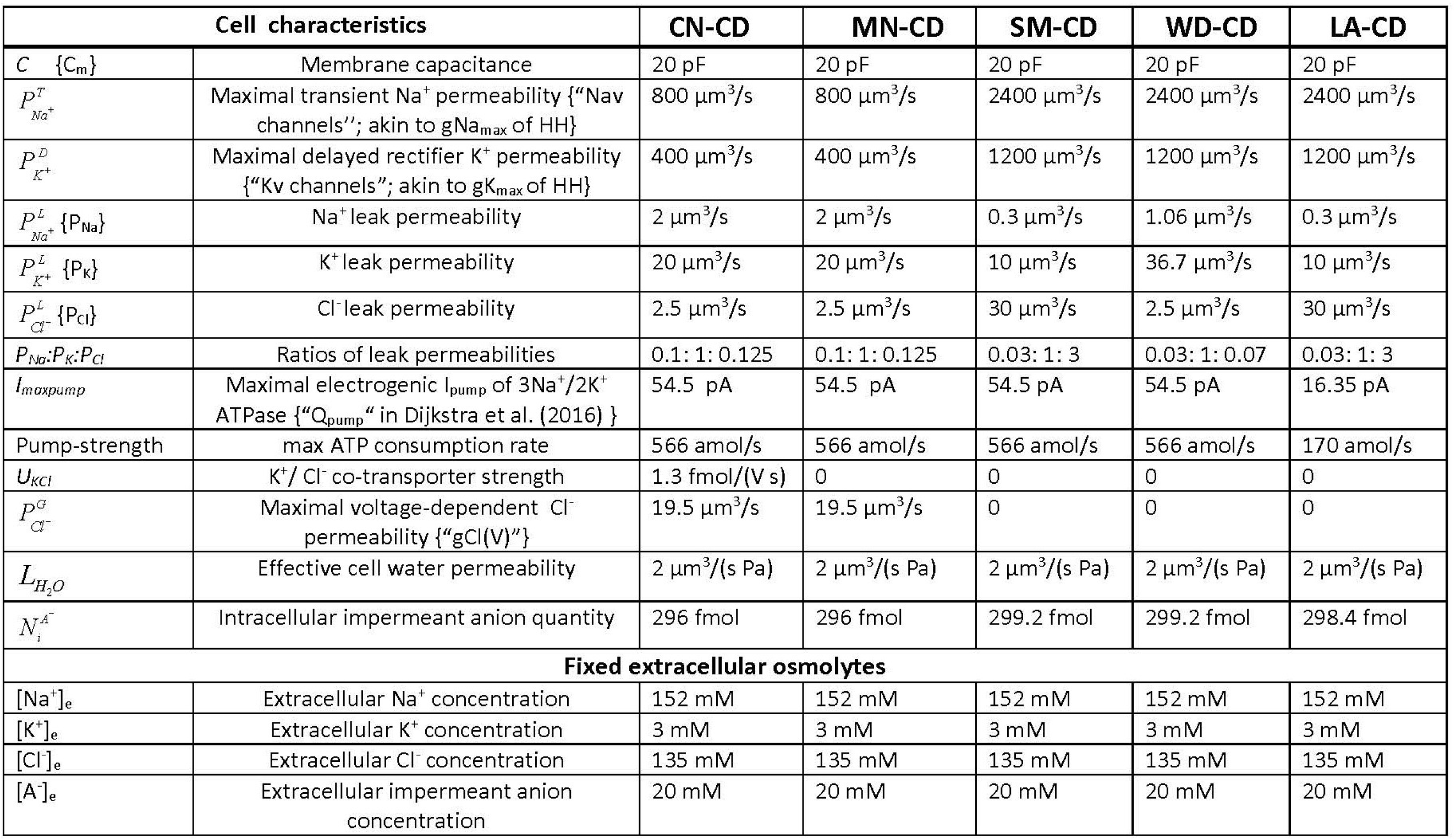
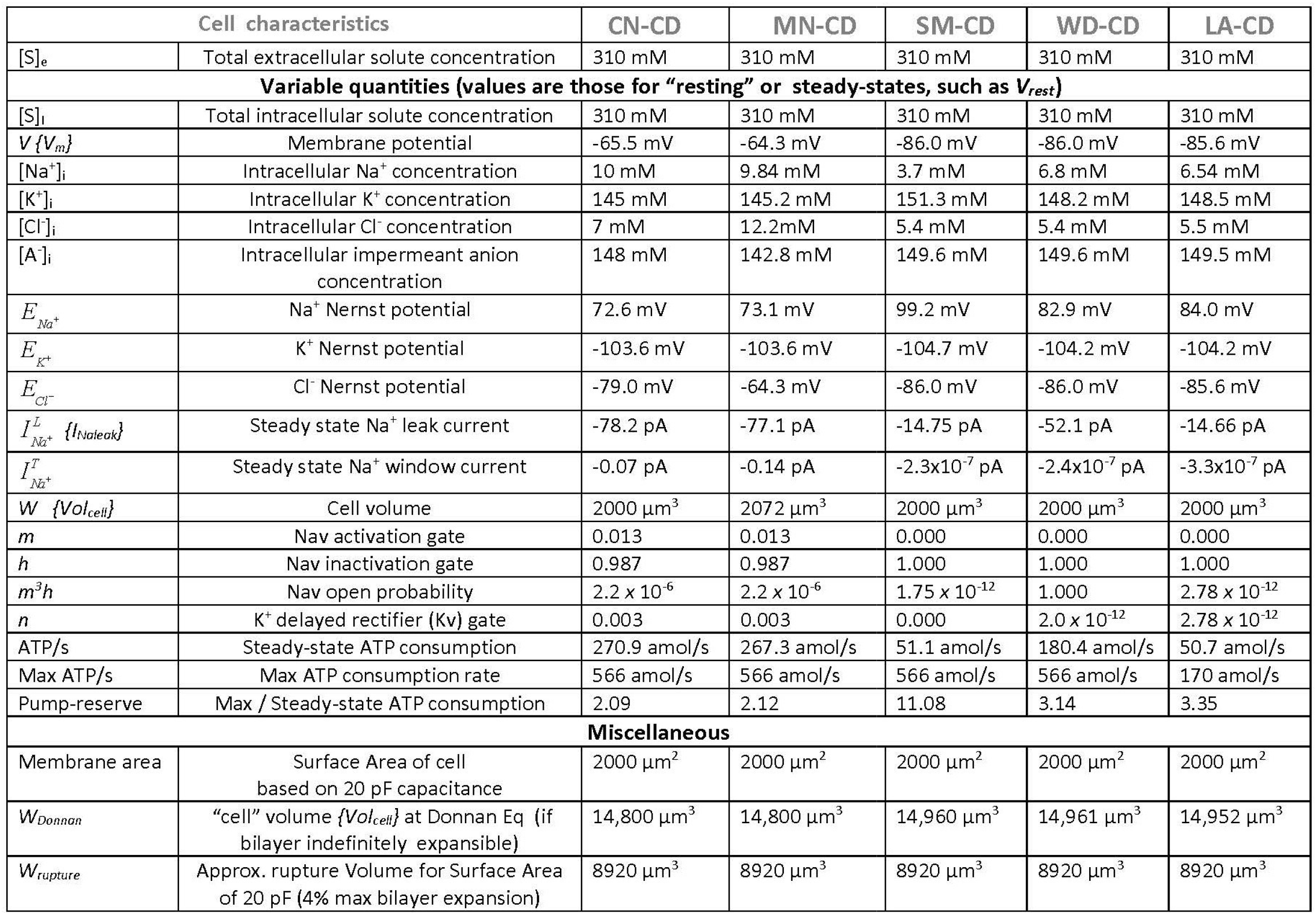
Parameter values for CN-CD and SM-CD models and variants. CN-CD entries are from Table 1 of Dijkstra et al (2016). For SM-CD values and other models, see Methods and text. When terms are listed two ways (e.g. *C* {C_m_}) the first is carried over from Dijkstra et al (2016) for use in equations but is usually referred to here (in text and figures) in the second way. All permeabilities (in units of μm^3^/s) are specific to the invariant cell area used here, as per Dijkstra et al (2016). (Constants: *F,* Faraday constant: 96485.333 C/mol; *R,* universal gas constant, 8.3144 J/K; *T,* Absolute temperature, 310 K; Specific membrane capacitance, 0.01 pF/μm^2^.)

| Cell characteristics |  | CN-CD | MN-CD | SM-CD | WD-CD | LA-CD |
| --- | --- | --- | --- | --- | --- | --- |
| $C \{C_m\}$ | Membrane capacitance | 20 pF | 20 pF | 20 pF | 20 pF | 20 pF |
| $P_{Na^+}^T$ | Maximal transient $\text{Na}^+$ permeability {"Nav channels"; akin to $g_{\text{Na}_{\text{max}}}$ of HH} | 800 $\mu\text{m}^3/\text{s}$ | 800 $\mu\text{m}^3/\text{s}$ | 2400 $\mu\text{m}^3/\text{s}$ | 2400 $\mu\text{m}^3/\text{s}$ | 2400 $\mu\text{m}^3/\text{s}$ |
| $P_{K^+}^D$ | Maximal delayed rectifier $\text{K}^+$ permeability {"Kv channels"; akin to $g_{\text{K}_{\text{max}}}$ of HH} | 400 $\mu\text{m}^3/\text{s}$ | 400 $\mu\text{m}^3/\text{s}$ | 1200 $\mu\text{m}^3/\text{s}$ | 1200 $\mu\text{m}^3/\text{s}$ | 1200 $\mu\text{m}^3/\text{s}$ |
| $P_{Na^+}^L \{P_{Na}\}$ | $\text{Na}^+$ leak permeability | 2 $\mu\text{m}^3/\text{s}$ | 2 $\mu\text{m}^3/\text{s}$ | 0.3 $\mu\text{m}^3/\text{s}$ | 1.06 $\mu\text{m}^3/\text{s}$ | 0.3 $\mu\text{m}^3/\text{s}$ |
| $P_{K^+}^L \{P_K\}$ | $\text{K}^+$ leak permeability | 20 $\mu\text{m}^3/\text{s}$ | 20 $\mu\text{m}^3/\text{s}$ | 10 $\mu\text{m}^3/\text{s}$ | 36.7 $\mu\text{m}^3/\text{s}$ | 10 $\mu\text{m}^3/\text{s}$ |
| $P_{Cl^-}^L \{P_{Cl}\}$ | $\text{Cl}^-$ leak permeability | 2.5 $\mu\text{m}^3/\text{s}$ | 2.5 $\mu\text{m}^3/\text{s}$ | 30 $\mu\text{m}^3/\text{s}$ | 2.5 $\mu\text{m}^3/\text{s}$ | 30 $\mu\text{m}^3/\text{s}$ |
| $P_{Na^+}P_{K^+}P_{Cl^-}$ | Ratios of leak permeabilities | 0.1 : 1 : 0.125 | 0.1 : 1 : 0.125 | 0.03 : 1 : 3 | 0.03 : 1 : 0.07 | 0.03 : 1 : 3 |
| $I_{\text{maxpump}}$ | Maximal electrogenic $I_{\text{pump}}$ of $3\text{Na}^+/2\text{K}^+$ ATPase {"$Q_{\text{pump}}$" in Dijkstra et al. (2016) } | 54.5 pA | 54.5 pA | 54.5 pA | 54.5 pA | 16.35 pA |
| Pump-strength | max ATP consumption rate | 566 amol/s | 566 amol/s | 566 amol/s | 566 amol/s | 170 amol/s |
| $U_{KCl}$ | $\text{K}^+/\text{Cl}^-$ co-transporter strength | 1.3 fmol/(V s) | 0 | 0 | 0 | 0 |
| $P_{Cl^-}^G$ | Maximal voltage-dependent $\text{Cl}^-$ permeability {"$g_{\text{Cl}}(V)$"} | 19.5 $\mu\text{m}^3/\text{s}$ | 19.5 $\mu\text{m}^3/\text{s}$ | 0 | 0 | 0 |
| $L_{H_2O}$ | Effective cell water permeability | 2 $\mu\text{m}^3/(\text{s Pa})$ | 2 $\mu\text{m}^3/(\text{s Pa})$ | 2 $\mu\text{m}^3/(\text{s Pa})$ | 2 $\mu\text{m}^3/(\text{s Pa})$ | 2 $\mu\text{m}^3/(\text{s Pa})$ |
| $N_i^A$ | Intracellular impermeant anion quantity | 296 fmol | 296 fmol | 299.2 fmol | 299.2 fmol | 298.4 fmol |
| Fixed extracellular osmolytes |  |  |  |  |  |  |
| $[\text{Na}^+]_e$ | Extracellular $\text{Na}^+$ concentration | 152 mM | 152 mM | 152 mM | 152 mM | 152 mM |
| $[\text{K}^+]_e$ | Extracellular $\text{K}^+$ concentration | 3 mM | 3 mM | 3 mM | 3 mM | 3 mM |
| $[\text{Cl}^-]_e$ | Extracellular $\text{Cl}^-$ concentration | 135 mM | 135 mM | 135 mM | 135 mM | 135 mM |
| $[\text{A}^-]_e$ | Extracellular impermeant anion concentration | 20 mM | 20 mM | 20 mM | 20 mM | 20 mM |
| [S]<sub>e</sub> | Total extracellular solute concentration | 310 mM | 310 mM | 310 mM | 310 mM | 310 mM |
| **Variable quantities (values are those for "resting" or steady-states, such as $V_{rest}$)** |  |  |  |  |  |  |
| [S]<sub>i</sub> | Total intracellular solute concentration | 310 mM | 310 mM | 310 mM | 310 mM | 310 mM |
| $V\{V_m\}$ | Membrane potential | -65.5 mV | -64.3 mV | -86.0 mV | -86.0 mV | -85.6 mV |
| [Na<sup>+</sup>]<sub>i</sub> | Intracellular Na<sup>+</sup> concentration | 10 mM | 9.84 mM | 3.7 mM | 6.8 mM | 6.54 mM |
| [K<sup>+</sup>]<sub>i</sub> | Intracellular K<sup>+</sup> concentration | 145 mM | 145.2 mM | 151.3 mM | 148.2 mM | 148.5 mM |
| [Cl<sup>-</sup>]<sub>i</sub> | Intracellular Cl<sup>-</sup> concentration | 7 mM | 12.2 mM | 5.4 mM | 5.4 mM | 5.5 mM |
| [A<sup>-</sup>]<sub>i</sub> | Intracellular impermeant anion concentration | 148 mM | 142.8 mM | 149.6 mM | 149.6 mM | 149.5 mM |
| $E_{Na^+}$ | Na<sup>+</sup> Nernst potential | 72.6 mV | 73.1 mV | 99.2 mV | 82.9 mV | 84.0 mV |
| $E_{K^+}$ | K<sup>+</sup> Nernst potential | -103.6 mV | -103.6 mV | -104.7 mV | -104.2 mV | -104.2 mV |
| $E_{Cl^-}$ | Cl<sup>-</sup> Nernst potential | -79.0 mV | -64.3 mV | -86.0 mV | -86.0 mV | -85.6 mV |
| $I_{Na^+}^L \{I_{NaLeak}\}$ | Steady state Na<sup>+</sup> leak current | -78.2 pA | -77.1 pA | -14.75 pA | -52.1 pA | -14.66 pA |
| $I_{Na^+}^T$ | Steady state Na<sup>+</sup> window current | -0.07 pA | -0.14 pA | -2.3x10<sup>-7</sup> pA | -2.4x10<sup>-7</sup> pA | -3.3x10<sup>-7</sup> pA |
| $W\{Vol_{cell}\}$ | Cell volume | 2000 μm<sup>3</sup> | 2072 μm<sup>3</sup> | 2000 μm<sup>3</sup> | 2000 μm<sup>3</sup> | 2000 μm<sup>3</sup> |
| $m$ | Nav activation gate | 0.013 | 0.013 | 0.000 | 0.000 | 0.000 |
| $h$ | Nav inactivation gate | 0.987 | 0.987 | 1.000 | 1.000 | 1.000 |
| $m^3h$ | Nav open probability | 2.2 x 10<sup>-6</sup> | 2.2 x 10<sup>-6</sup> | 1.75 x 10<sup>-12</sup> | 1.000 | 2.78 x 10<sup>-12</sup> |
| $n$ | K<sup>+</sup> delayed rectifier (Kv) gate | 0.003 | 0.003 | 0.000 | 2.0 x 10<sup>-12</sup> | 2.78 x 10<sup>-12</sup> |
| ATP/s | Steady-state ATP consumption | 270.9 amol/s | 267.3 amol/s | 51.1 amol/s | 180.4 amol/s | 50.7 amol/s |
| Max ATP/s | Max ATP consumption rate | 566 amol/s | 566 amol/s | 566 amol/s | 566 amol/s | 170 amol/s |
| Pump-reserve | Max / Steady-state ATP consumption | 2.09 | 2.12 | 11.08 | 3.14 | 3.35 |
| **Miscellaneous** |  |  |  |  |  |  |
| Membrane area | Surface Area of cell based on 20 pF capacitance | 2000 μm<sup>2</sup> | 2000 μm<sup>2</sup> | 2000 μm<sup>2</sup> | 2000 μm<sup>2</sup> | 2000 μm<sup>2</sup> |
| $W_{Donnan}$ | "cell" volume {$Vol_{cell}$} at Donnan Eq (if bilayer indefinitely expansible) | 14,800 μm<sup>3</sup> | 14,800 μm<sup>3</sup> | 14,960 μm<sup>3</sup> | 14,961 μm<sup>3</sup> | 14,952 μm<sup>3</sup> |
| $W_{rupture}$ | Approx. rupture Volume for Surface Area of 20 pF (4% max bilayer expansion) | 8920 μm<sup>3</sup> | 8920 μm<sup>3</sup> | 8920 μm<sup>3</sup> | 8920 μm<sup>3</sup> | 8920 μm<sup>3</sup> |

SM-CD P-L/D resilience substantially exceeds that of CN-CD, a differential even more impressive if SMFs’ distinctive syncytial morphology is factored in. Joos and Morris (submitted) estimate that, *in situ,* 2000 μm^2^ would service a myoplasmic volume ~10-15X that of SM-CD. Steady-state P-L/D values depend only on surface area, syncytial morphology therefore magnifies muscle tissue’s resting efficiency. SMFs’ rates of change (e.g., during rundown and recovery) would be slower than depicted by SM-CD, but for rundown, this is a boon, while for recovery, given SMFs’ large pump-reserve and low duty-cycle, it would generally be unproblematic.

### Small P_Na_, pump-reserve, *mdx* fibers, a chronically ischemic fiber model

Hyperpolarizing I_pump_ counter-balances depolarizing I_Naleak_ across P_Na_ in P-L/D systems; maximal hyperpolaring I_pump_ in SM-CD and CN-CD are identical. Though its P_K_:P_Na_ ratio makes resting SM-CD ~20 mV more negative than resting CN-CD, [small I_Naleak_] makes SM-CD’s ATP-consumption 5.3X lower (its pump-reserve is thus 5.3X larger; pump-reserve is maximal/resting ATP-consumption or its equivalent, maximal/resting I_pump_)(**Table 2**). Small P_Na_ is thus pivotal for SMFs’ extreme ion homeostatic efficiency and for their pump-reserve based robustness. Pump-reserve is 11-fold in SM-CD, while for rat muscle, 7-fold to 22-fold is reported (see Clausen 2013, 2015).

Pump-protein density in isolated *mdx* sarcolemma (Anderson 1981; Dunn 1993) exceeds that of control mice. Nevertheless, we assume that membrane-damage etc (**Figure 1A**) puts *<u>operational</u> mdx* pump-strength in the “ischemic” range (<100% of normal). In *mdx* fibers NF-κB-dependent modulators depress pumping and increase the Na^+^-overload of (**Table 1** item **4**; Miles at al 2011). In *mdx* fibers, stimulating NO pathways almost fully abolishes the Na^+^-overload of (Altamirano et al 2014; **Table 1**, item **7**); whether (as presumed) this is attributable to “more Na^+^-pumping” was not determined. In a rat diaphragm DMD-model (Lehmann-Horn et al 2012), eplerenone up-regulation of the Na/K-ATPase (via Tyr10 dephosphorylation of its α-subunit) results in ouabain-sensitive fiber repolarization (Breitenbach et al 2016). Already prescribed for DMD cardiomyopathy (Raman et al 2017), eplerenone has been considered for DMD skeletal muscle (pilot study; Glemser et al 2017; **Table 1**, items **5**,**6**,**9**).

While enabling rapid post-perturbation return to steady-state, large pump-reserves also provide generous capacity for ENa-exploiting Na^+^-transporters. Since SM-CD lacks Na^+^-transporters, its steadystate [Na^+^]_I_ (3.7 mM) is understandably lower than in healthy mouse SMFs (**Table 1** items **2,4,7,8**; i.e., 5.3-13 mM). Including transporters would complicate attempt to assess whether chronic Na^+^-overload could develop solely from: **1**) too little Na^+^-pumping (ischemia) or **2**) too much Na^+^-leaking (leaky channels). Using *mdx*-variants, Burr et al (2014), showed that reverse operation of Na^+^/Ca^2+^-exchangers can trigger Ca^2+^-necrosis in SMFs, provided there is a pre-existing Na^+^-overload. To design therapeutic approaches, mechanistic clarity on the provenance of Na^+^-overload is needed.

To represent deeply ischemic viable DMD fibers, LA-CD (30% pump-strength of SM-CD, pump-reserve=3.4-fold, (**Table 2**)) is used. Its 6.5 mM [Na^+^]_i_ is still within the range reported for healthy mice but if, as in working mouse fibers, Na^+^-transporters (e.g., Na^+^/H^+^ exchanger; Iwata et al 2007) were depleting E_Na_, [Na^4^]_I_ could easily rise into the (chronically Na^+^-overloaded) range reported for *mdx* fibers by Burr et al (2014) (**Table 1**, item **8**; 7.3 mM).

Even ignoring that healthy rodent SMF pump-reserves far exceed the 3.4-fold of LA-CD, it ion homeostasis is sub-par based on this: to maintain [Na^+^]_i_=6.5 mM, LA-CD consumes ATP at 50.7 amol/s, whereas SM-CD, consuming essentially the same amount (51.1 amol/s), achieves [Na^+^]_i_=3.7 mM (see **Figure 9 Bii**).

### Small input impedance and [big P_Cl_] in ischemic SM-CD

Small P_Na_ is imperative for the low-cost steady-state that lets chronically ischemic DMD fibers survive. For strongly hyperpolarized SMFs, [small I_Naleak_] through small P_Na_ is workable because, to keep their excitability suitably low, SMFs use *not* big P_K_, but [big P_Cl_]. WD-CD, a counterfactual Pump-Leak dominated analog of SM-CD (same low input impedance, same V_rest_ but “big P_K_, big P_Na_, small P_Cl_”; **Table 2**) shows this quantitatively (Joos & Morris, submitted).

In *mdx* (Cozzoli et al 2014) as in healthy fibers open ClC-1 channels dominate [big P_Cl_]. Unwanted firing is detrimental to ischemic DMD fibers. To supress such firing without metabolic penalty would be attractive. Excitability modulation via ΔP_Cl_ (Pedersen et al 2011, 2016) therefor reveals an ancillary benefit of the Donnan dominated strategy: in SM-CD, changing P_Cl_ say, +10-fold changes neither steady-state ATP-consumption (51.1 amol/s) nor V_rest_ (−86.0 mV). Likewise for LA-CD. Electrophysiologically, this accords, e.g., with the finding that SMFs from wild-type and ClC-1-gene-null mice have the same V_rest_ (Novak et al 2015). Further, Bækgaard Nielsen et al (2017) find that, with heavy exercise, fast-twitch fibers initially inhibit ClC-1s but as stimulation continues, inhibition can suddenly reverse to dramatically increased ClC-1-opening, abolishing excitability.

The SMF/neuron contrast is especially stark here: whereas ↑P_Cl_ hastens cytotoxic swelling in ischemic neurons (Rungta et al 2015; Dijkstra et al 2016; Joos and Morris submitted), ischemic SMFs deploy ↑P_Cl_ for enhanced safety.

### Small P__Na__: anoxic rundown and recovery in SM-CD and LA-CD

Transient functional ischemia imperils DMD fibers. In **Figure 2Ai**, with I_pump_=0, anoxic rundown of SM-CD to Donnan Equilibrium (DE) is slow (as per Joos & Morris submitted) but here, in addition, we examine pump-strength restoration (to 100%). Restoring pumps during rundown or even during ectopic firing (**Figure 2Aii,iii**) lets the system return quickly to steady-state; [Na^+^]_I_-sensitive I_pump_ hyperpolarizes Vm until [Na^+^]_I_ renormalizes. Critically, however, once ectopic firing stops, pump-restoration no longer supports recovery (**Figure 2Aiv**); degradation continues, not to Donnan Equilibrium (DE), but to towards a pathological steady-state, DE-like, though ATP-consuming.

**Figure 2.**
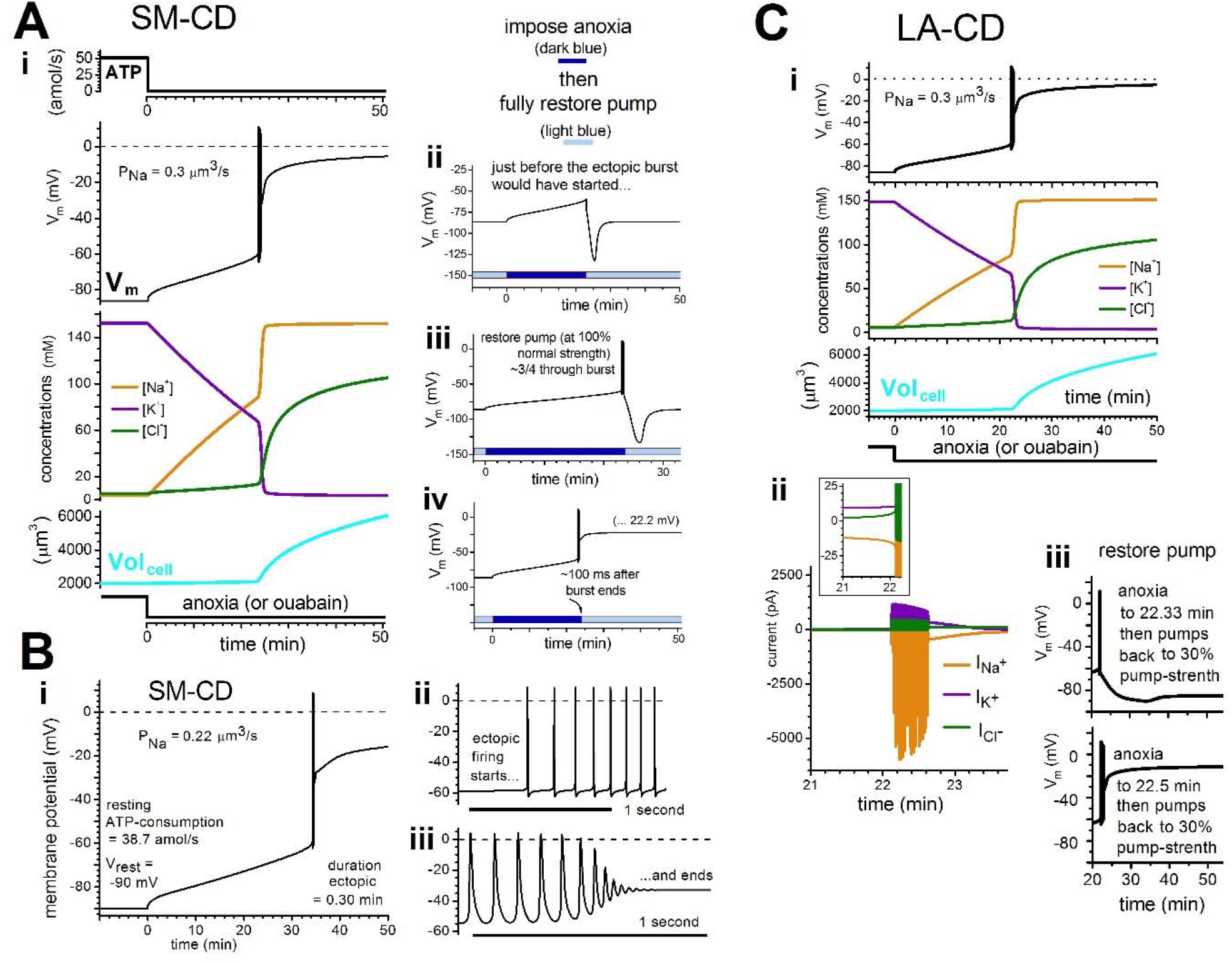
Anoxic rundown and recovery. **Ai.** anoxic rundown of SM-CD (experimental equivalent: instantaneous exposure to pump-inhibitor, ouabain). Following the small pump-off depolarization, rundown is limited by [small I_Naleak_]. Since P_K_>>P_Na_, concurrent K^+^ efflux can almost maintain cytoplasmic electroneutrality. With P_Cl_ so large, ΔE_Cl_ tracks ΔV_m_, minimizing the driving force on Cl^-^; rundown-swelling is minor because, with Δ[Na^+^]_i_ rate-limiting, ↑[Na^+^+Cl^-^]_i_ is minor. Finally, 22.7 minutes into anoxia, SM-CD reaches the Nav channel activation threshold; I_Na_ from “myogenic” APs elicits a catastrophic concurrent [Cl^-^+H_2_O] influx. SM-CD swells and degrades toward DE. **ii, iii**, **iv,** SM-CD Vm(t) before/during/after anoxia is imposed, with 100% pump-strength restored at different times, as labeled**. Bi.** In SM-CD a ↓P_Na_ (as labeled and explained in the text) prolongs the anoxic rundown time to ectopic firing, whose start and termination is expanded in **ii, iii**. **Ci,** anoxic rundown of LA-CD (as per **Table 2**, steady-state Volcell for LA-CD and SM-CD are identical because the impermeant cytoplasmic anion quantity in LA-CD was slightly decreased). **ii,** ion currents in the minutes before/during/after spontaneous myogenic firing (the inset box’s greatly expanded vertical resolution emphasizes the rundown currents before firing), **iii,** restoring LA-CD pump-strength (30%) during myogenic firing (top) and just after it ceases (bottom).

Nevertheless, even with pumps-off, note that the “[big P_Cl_][small I_Naleak_]” “architecture” of SMF ion homeostasis offers exceptional protection in the form of a prolonged period of reversible rundown (issues relating to ischemia-reperfusion injury (Dudley et al 2006; Schmucker et al 2015; Li et al 2020) are beyond our scope). The [small I_Naleak_] is rate-limiting and keeps fully-anoxic SM-CD safe for 22.7 minutes. During the rundown, E_Na_ and E_K_ depletion is almost matched, and the extent of swelling is modest. Given the SA/V ratios of actual myofibers (as explained above) their “safe” anoxic-rundown durations would substantially exceed SM-CD’s.

**Figure 2Bi** shows that a slight reduction of already-small P_Na_ markedly increases its benefits. For SM-CD, V_rest_=-86 mV. When P_Na_ drops from 0.3 μm^3^/s→0.22 μm^3^/s, V_rest_=-90 mV and ATP-consumption falls (51.1 amol/s→38.7 amol/s), boosting pump-reserve from 11.1-fold→14.6-fold. With P_Na_ reduced, anoxic rundown slows: ectopic firing begins at 34 not 22.7 minutes (**2Bii,iii** expands the start and termination of firing) while recovery initiated during rundown (not shown) is quicker.

The primacy of small P_Na_ is made further evident by the fact that anoxic rundown speed for SM-CD is almost indifferent to P_Cl_: SM-CD “goes ectopic” at 20.7 minutes for 0.1XP_Cl_, and at 23.8 minutes for 10XP_Cl_ (not shown). Thus, while ↑P_Cl_ could abruptly ↓endplate-triggered excitability under energy-depleted conditions (Bækgaard Nielsen et al 2017).e.g., ↑P_Cl_ would not appreciably delay the onset of myogenic firing in ischemic (DMD) fibers. By contrast, in such fibers, ↓P_Na_ would hyperpolarize Vm, thereby lowering endplate excitability, and would also delay myogenic firing.

Deeply ischemic LA-CD has the same P_Na_ as SM-CD but its higher [Na^+^]_I_ and slightly depolarized V_re_st (**Table 2**) slightly reduce its Na^+^ driving force. As for SM-CD, the anoxic rundown pace of LA-CD is P_Na_-limited. As seen in **Figure 2Ci**, ectopic firing commences in LA-CD at 22.0 minutes (22.7 for SM-CD). During ectopic firing LA-CD’s AP-associated currents (**Figure 2Cii**) differ inconsequentially from those (not shown) of SM-CD. Nevertheless, LA-CD can recover provided its pumps (30%) are restored before ectopic firing ends (**Figure 2Ciii**); as expected, recovery is slower than for SM-CD (which gets 0%→100%, not 0%→30%). Consistent with their syncytial SMF geometry, chronically-ischemic DMD fibers recovering after a comparable bout of transient functional ischemia/anoxia would exhibit considerably more prolonged recovery periods than LA-CD.

### “Too little Na^+^-pumping” alone could explain chronic Na^+^-overload in DMD

While LA-CD’s elevated [Na^+^]_I_ already hints that ischemia alone (“too little Na^+^-pumping”) might suffice to explain *mdx-*level Na^+^-overloads, pump-strength effects need to be examined systematically.

Excitable P-L/D systems have non-linear driving forces and two interacting non-linear Na^+^ fluxes (I_pump_([Na^+^]_i_) and I_Nav_(V_m_)). Even for ultra-simple SM-CD, therefore, predicting/describing how ion homeostasis plays out across pump-strengths requires graphical analysis. Joos & Morris (submitted) showed that across its full pump-strength range (up-regulated several-fold down to anoxic), SM-CD, like other excitable P-L/D systems (e.g., CN-CD; Dijkstra et al 2016; Hübel et al 2014), exhibits bistability, i.e., discontinuous physiological and pathological steady-state continua accessed via two unstable thresholds (see **Figure 3A**).

**Figure 3.**
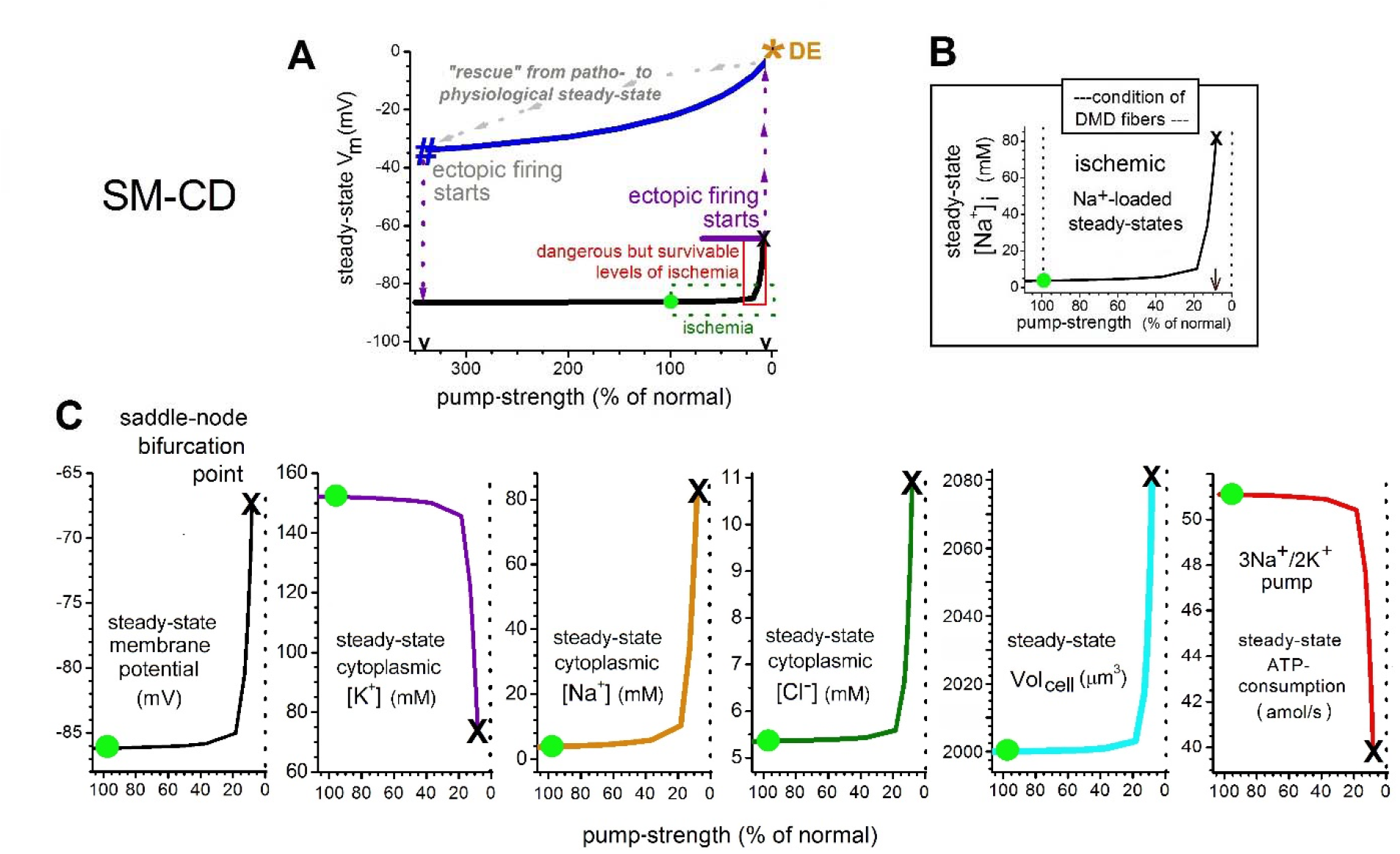
SM-CD steady-state bifurcation plots. A. The SM-CD bifurcation plot for steady-state V_m_ as a function of pump-strength; the steady-state or stable values form continua; unstable values occur at points **#** and the saddle-node, **X**; DE is the Donnan Equilibirum point). This illustrates what it means to say that steady-state P-L/D values vary non-linearly (and bistably) as pump-strength varies (see Joos and Morris, submitted); for DMD fibers, the ischemic “danger zone” (boxed region), which includes the saddle-node (**X**), is critical. **B.** Ischemic range for steady-state [Na^+^]_i_ up to the unstable saddle node. **C**. Ischemic region for each of the SM-CD parameters, as labelled. Approaching the saddle-node, i.e., in the danger zone, the system shows a steep IATP-consumption (=↓hyperpolarizing I_pump_) concomitant with a steep ↑[Na^+^]_i_ (~10 mM to ~90 mM). This occurs because from 10-90 mM, pump-rate sensitivity to [Na^+^]_i_ (see **Fig methpump**) is radically shallower than close to normal [Na^+^]_i_ (i.e. up to ~10 mM). In the danger zone, an I_pump_ value sufficient to achieve steady-state requires a large [Na^+^]_I_, but this entails E_Na_ depletion and concomitant E_K_ depletion. The resultant depolarization further diminishes I_Naleak_; hence the ↓ATP-consumption (at steady-state, recall, -I_Naleak_=hyperpolarizing I_pump_). Finally, when pump-strength is so low that pump-reserve<1, the excitable P-L/D system is depolarized to **X**, i.e. to its unstable saddle-node.

For chronically ischemic DMD however, only the ischemic-but-viable physiological steady-states (**Figure 3B,C**) are relevant. For V_m_(pump-strength) (**Figure 3A**) only, the full bifurcation plot is provided to give context for partial plots **Figure 3B,C**. Beyond the saddle-node(**X**), ectopic firing induced Cabnecrosis would render the pathological steady-state continuum (**Figure 3A**) irrelevant for DMD (in *mdx,* ectopic Ca^2+^-mediated contractures destroy the fibers (Claflin and Brooks, 2008)).

For DMD patients,[Na^+^]_I_ is a measurable ion homeostatic parameter, so **Figure 3B** emphasizes the physiological steady-state continuum for [Na^+^]_I_(pump-strength) in the ischemic (<100%) range. On these plots, our DMD mimic, LA-CD, essentially corresponds to 30%. Thanks to its [small I_Naleak_], ischemic SM-CD maintains pump-reserve>1-fold until pump-strength drops to ~8%, below which (i.e. at its saddle-node(**X**)), the system destabilizes. If (as per **Figure 2A→B)** P_Na_ were to decrease, would hold off to an even lower pump-strength. SM-CD could be considered “chronically Na^+^-overloaded” from ~20-15% down to 8%. Thus, P-L/D analysis indicates that, in principle, DMD/*mdx c*hronic Na^+^-overload could arise solely from “too-little-Na^+^-pumping.”

When severe ischemia subsides, recovery by Donnan dominated SM-CD is an incomparably more robust than for Pump-Leak dominated neurons (Joos & Morris, submitted) where the saddle-node is at 65% pump-strength; if pump-strength falls to, say, 64%, only a pump-strength increase >181% allows for recovery (Dijkstra et al 2016). For SM-CD, however, any ↑blood-flow (i.e., any ↑pump-strength to a level >8%) contributes to recovery.

For DMD ischemia, therapeutic avenues for ↑pump-strength include counteracting vascular insufficiency (e.g., angiogenesis (Podkalicka et al 2019), ↑vasodilation (e.g.,NO-reagents (Patel et al 2018)) and ↑functional pump-density (Breitenbach et al 2016; Glesmer et al 2017; Raman et al 2017).

### The danger zone for DMD

For physiological SM-CD steady-states in the ischemic range, **Figure 3C** assembles the P-L/D parameter plots. This shows that above ~40% pump-strength, SM-CD is extremely robust, while below ~20%, system deterioration is steep. In this “danger zone” Vm depolarizes appreciably, while [Na^+^]_i_ and [Cl^-^] rise and [K^+^]_i_ falls precipitously (note absolute amounts). Because P_K_>>P_Na_, concurrent Na^+^-loading/K^+^-depletion largely preserve cytoplasmic electroneutrality. Even at the saddle-node, where SM-CD [Na^+^]_I_=88 mM, Cl^-^-loading is so minimal that swelling is only ~4.5% (2000μm^3^→~2090μm^3^). As Vol_cell_ increases, impermeant anions dilute slightly (149.6mM→143.1mM for 100%→saddle-node; not shown). Thus, in spite of [big P_Cl_], chronic ischemia-induced Na^+^-overloads are almost “non-osmotic”. This accords with recent MRI findings from DMD patients’ whose muscles show chronic Na^+^-overload even in the absence of “water T2 alterations” (Gerhalter et al 2019). Any demonstrably-swollen Na^+^-overloaded fibers in patients would likely be experiencing myogenic APs.

Though computational ATP is never limiting in our analyses, note the situation below 20% pump-strength; as [Na^+^]_I_(pump-strength) rises steeply, ATP-consumption declines abruptly. As detailed further in the **Figure 3** legend, this can equivalently be described as a steep decline in hyperpolarizing I_pump_ (=-I_Naleak_) in this zone; briefly, falling ATP-consumption reflects pump saturation, ENa dissipation and specifics of the P-L/D system’s I_pump_/[Na^+^]_i_ feed-back interactions.

P-L/D bifurcation analysis of SM-CD thus shows that deep ischemia (“too little Na^+^-pumping”) yields chronic “non-osmotic” Na^+^-overload at levels that are tolerated because the Donnan dominated system’s “control valve” (its extremely small P_Na_) has facilitated its large pump-reserve.

This in no way implies that damage-induced Na^+^-leaks do not also contribute to DMD Na^+^-overload, only that ischemia, if sufficiently deep, would suffice. For “milder” ischemia (say, pump-strengths>25%) chronic Na^+^-overload would (by elimination) also reflect Na^+^-leaks and/or over-active Na^+^-transporters. Bleb-damage sections below address leaky Na^+^-permeant channels.

The concentration of extracellular Donnan effector is invariant here. However, as Mehta et al (2008) indicate, for very small pump-reserve, this parameter would affect steady-state. In the danger zone, pump-reserve is increasingly marginal. Since Coles et al (2020) report protease-release of extracellular matrix proteins in exercised *mdx* fibers, extracellular Donnan effector impacts (influenced, probably, by ischemia-related ΔpH (Hagsberg 1983)) could be worth revisiting.

### Sleep, diaphragm fibers and the danger zone

Though DMD patients avoid over-exertion, even their sleep holds dangers (Hartman et al 2020). Skeletal muscle tissue is no known to be directly involved (Ehlen et al 2017) in generating sleep rhythms, including variations in SMF glucose homeostasis and oxidative capacity (Harfmann et al 2016; van Moorsel et al 2016). During REM sleep, DMD patients’ impaired sleep and respiratory profiles become acutely compromised (Nozoe et al 2015). For already-ischemic DMD diaphragm fibers, diminished energy status during sleep could take them treacherously far into the danger zone. For healthy steady-states, pump-model specifics matter little (Düsterwald et al 2018). However, for deeply ischemic excitable cells, pump-specifics that affect a P-L/D danger zone’s inflection region and steepness could be important. Future SM-CD refinements focusing on the DMD danger zone might include using SMF-specific pump model, perhaps incorporating dystrophic fibers’ pump characteristics (Hakimjavadi et al 2018; Kravtsova et al 2020). Similarly, since V_rest_ is chronically depolarized in the danger zone, Nav gating particulars (e.g., Filatov et al 2005) could be added to accurately depict fiber excitability.

### DMD bleb-damage: “too much Na^+^-leak”

In SMFs, P_K_>>P_Na_. Therefore, the major consequence of any “leaky cation channel” will be augmented P_Na_. Specifically, in SM-CD and LA-CD, P_K_:P_Na_ is 1.00:0.03 so if an SMF cation channel with, say, P_K_:P_Na_=1:1 over-activated enough to double P_Na_ (100% increase), P_K_ would increase only 3%. This is relevant because in many excitable cells, a hormonally-regulated cation channel, NALCN (**NA L**eak **C**hannel **N**on-selective) contributes to resting P_Na_ (Lu et al 2007; Cochet-Bissuel et al 2104; Lutas et al 2016; Philippart and Khaliq, 2018; Reinl et al 2018). No NALCN is detected in SMFs, but other cation channels (Metzger et al 2020) might contribute to P_Na_ and if over-active, they would be problematic. In *mdx* fibers, over-active mechanosensitive cation channels (unidentified) and AChRs are posited routes for pathological Ca^2+^-entry (Franco and Lansman 1990; Carlson and Officer 1996; Yeung et al 2005; Lansman 2015; Ward et al 2018). If so, unavoidably, they are also “Na^+^-leaks”. The mechanosensitive channel is a suggested DMD therapeutic target for GsMtx4-related peptides (Ward et al 2018; Gnanasambandam et al 2017).

The unspecified computational “leaky cation channel” modeled here could represent either, both, or neither of these cation channels but it is assigned P_K_:P_Na_=1.11:1, the ratio of SMF AChRs (Hille 2001). AChRs are abundant, identified, and implicated as leaky in *mdx* (Carlson and Officer 1996). They are also mechanosensitive, GsMtx4-inhibited (Pan et al 2012). Plausibly, their spontaneous activation (Jackson et al 1990) might increase in damaged junctional *mdx* fiber sarcolemma (Pratt et al 2015).

Healthy or dystrophic, SMFs’ biggest Na^+^ influxes are via Nav1.4 channels (Fu et al 2011). Nav1.4 binds syntrophin and thence dystrophin (whether utrophin can substitute is unknown (Gee et al 1998; Albesa et al 2011)). Sarcolemmal Nav1.4 density is somewhat low in *mdx* (Ribaux et al 2001; Hirn et al 2008) but Nav1.4 gating appears normal (Mathes et al 1991). Hirn et al (2008) showed that for 3 days after mechanically-traumatic fiber isolation, 3 nM tetrodotoxin protects *mdx* fibers from Na^+^-loading and dieoff (**Figure 1B**). In DMD sarcolemma, Na^+^-leak through Nav1.4 channels would likely take the form of left-shifted window-current from Nav channels in bleb-damaged sarcolemma (see “Nav-CLS”, **Figure 1D;** Morris et al 2012a, b). The clinical Nav-antagonist, ranolazine, is a proposed DMD therapeutic (Burr et al 2014; Lorusso et al 2019).

The next sections show SM-CD (and other models) dealing with both classes of Na^+^-leak (Nav-CLS, over-active cation channels).

### Bleb-damage: Nav-CLS is treacherous but cannot explain chronic Na^+^-overload

**Figure 4** shows P-L/D responses to increasingly intense Nav-CLS (**Figure 1D** and **Methods**) from bleb-damage to a fraction (0.3) of the Nav-bearing sarcolemma. As detailed in the figure legend, CN-CD (**Figure 4A**) and SM-CD (**Figure 4B**) are compared. Since Pump-strength=100%, P-L/D feedback is initially able to counteract SM-CD’s damage-induced ectopic firing. This differs from ectopic firing due to anoxia (**Figure 2A**) or to extreme-ischemia (Joos & Morris submitted, their Fig 7Aii) which inexorably ends in cytotoxic swelling. Until the system destabilizes at Nav-CLS(0.3)=~30 mV, this Nav-CLS-induced ectopic firing causes minimal swelling. The resultant ectopic contractures (e.g. at Nav-CLS(0.3)=28 mV; Figure **4Bii**) would, however, be very problematic *in situ.*

**Figure 4.**
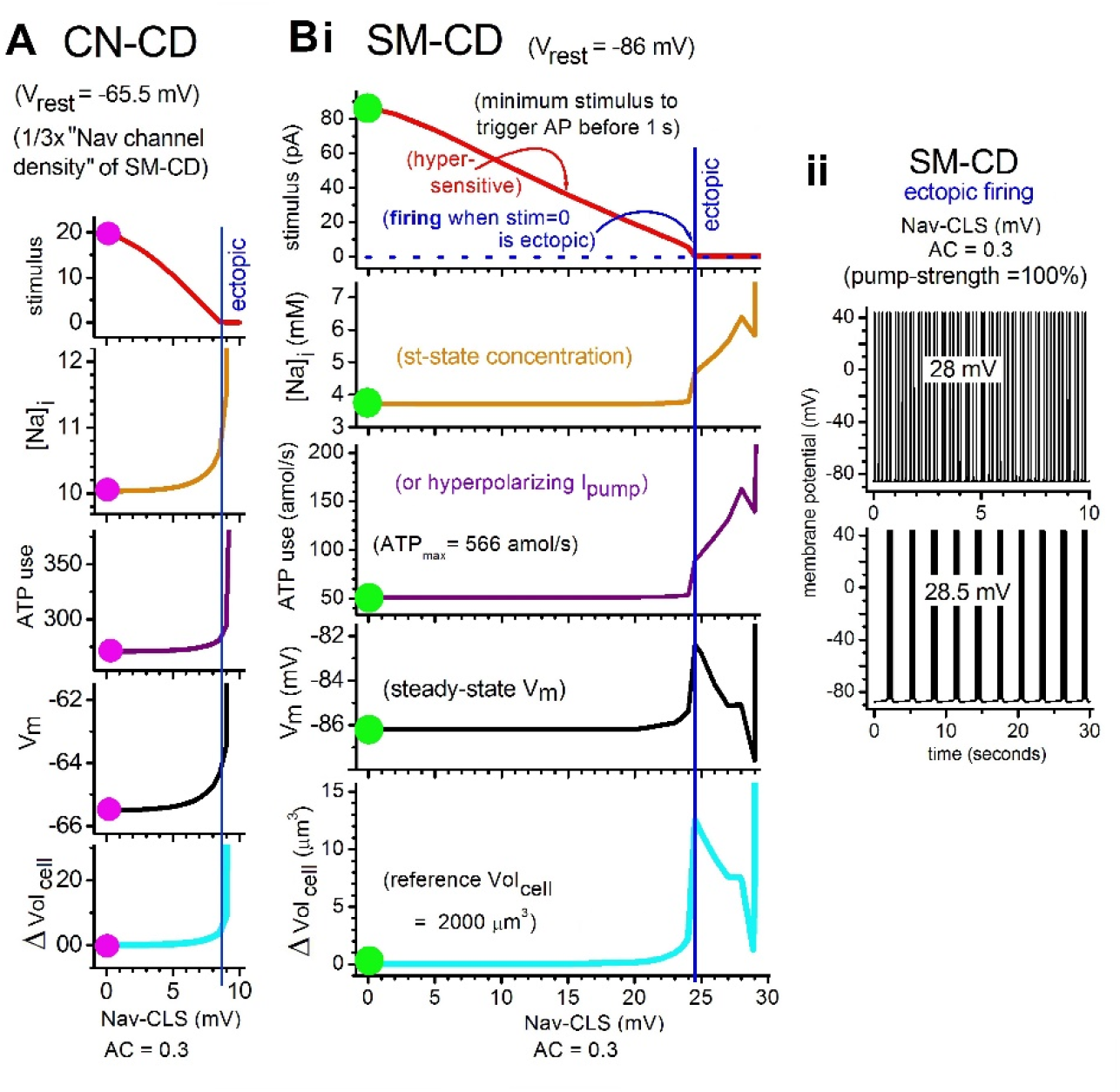
Bleb-damage induced Nav-CLS in CN-CD and SM-CD. For CN-CD (**A**) and SM-CD (**B**), 30% (affected channels, i.e., AC = 0.3) of the Nav-bearing membrane is damaged at Nav-CLS intensities ranging from 0 mV (magenta and green dots = healthy steady-state) up to 10 mV for CN-CD or up to 30 mV for SM-CD. Most (70%) of the Nav channels are as depicted in **Figure 3B** but 30% (the Navs in bleb-damaged membrane), having undergone Nav-CLS, have their m^3^ and h shifted by the indicated amount (the non-dimensional “window conductance” consequence of NavCLS=20 mV is illustrated in **Figure 1D**). **Bii**, for SM-CD, Vm(t) as labeled. **A** and **B**, top panels reveal subthreshold Nav-CLS effects; negative slopes indicate increasing hypersensitivity to a test stimulus. Vertical blue lines indicate where spontaneous “ectopic” (or myogenic for SMFs) firing begins. Beyond that, CN-CD’s small pump-reserve cannot counteract the Na^+^-loading; [Na^+^+Cl^-^+H_2_O]_i_ (thence Vol_cell_) increases catastrophically (=cytotoxic swelling). This accords with excitation-related Na^+^-overload computations of Dijkstra et al (2016) but shows more biophysical nuance because Nav-CLS intensity is graded and Nav-CLS onset is instantaneous (to mimic excitotoxic veratridine, Dijkstra et al set the H-H parameter, *h,* to 1, and provided a “wash-in” period). Its strongly hyperpolarized V_rest_ lets SM-CD withstand more intense Nav-CLS damage (than CN-CD) before firing ectopically. Thanks to SM-CD’s big pump-reserve (pump-strength= 100% here), firing, when first it develops, does not elicit cytotoxic swelling.). At 28 mV it causes somewhat irregular 5-6 Hz tonic firing and 28.5 it causes bursts of 16-18 APs, with intra-burst frequency ~25 Hz. Only at Nav-CLS=30 mV, does SM-CD succumb to swelling. The Cl^-^-entry plot (not shown) has the same shape as Volcell(Nav-CLS). Beyond the blue line (Nav-CLS-induced myogenic firing), SM-CD’s complex parameter changes reflect P-L/D feedback plus the changing excitation patterns. As per Boucher et al (2012), tonic firing gives way to bursts (**Bii**) then (not shown) to depolarizing block. At Nav-CLS=28 mV and 28.5 mV, a ~3-fold ↑ATP-consumption prevents major Na^+^-loading ([Na^+^]_i_ reaches only ~6 mM). Swelling is minimal. At Nav-CLS=~29 mV, Ipump hyperpolarizes V_rest_. At 30 mV, with (hyperpolarizing Ipump)<(depolarizing (I_Naleak_+ time-averaged INav)), the P-L/D system is overwhelmed.

Other DMD patho-phenomena here include subthreshold hypersensitivity (**4Bi** top panel) that would foster myogenic APs. This likely contributes to DMD’s erratic spontaneous contractility (Ishpekova at al 1999; Nojszewska et al 2017). Single fibre EMG recordings from advanced-stage DMD patients show bizarre repetitive discharge bursts (Trontelj and Stålberg, 1983; see Yu et al 2012). In *mdx* muscles, electromyography of abnormal spontaneous potentials and complex repetitive discharges resemble the muscle discharges of boys with DMD (Carter et al 1992; Han et al 2006). Where there is Nav-CLS-induced hypersensitivity, erratic myogenic APs bouts are expected if the damaged dystrophic sarcolemma also has leaky” cation-channels that activate intermittently, depolarizing Vm (Lansman 2015; and see next section).

The exquisitely narrow Nav-CLS damage range between zero Na^+^-loading and catastrophic instability ([Na^+^]_i_ panel, **Figure 4Bi**) makes Nav-leak an implausible stand-alone explanation for DMD’s chronic Na^+^-overload. Nevertheless, the considerable danger posed by subthreshold Nav-CLS effects (hypersensitivity) and myogenic firing make it important to determine if slow-inactivation of Nav1.4 (Webb et al 2009; Morris et al 2012A,B; Lukacs et al 2018) protects SMFs during the membrane repair/regeneration processes critical for DMD longevity (Baumann et al 2020). However, even for otherwise healthy fibers recovering from overexertion-tears (Dahlmann et al 2016; Hotfiel et al 2018; Hody et al 2019) this seems not to have been examined.

### Bleb-damage: depolarizing and Na^+^-loading effects of leaky cation channels

In **Figure 5** cation-channels are activated (P_Na_-component increased by 0.125 μm^3^/s; see legend and **Methods**) for 4 minutes in SM-CD, LA-CD, and in WD-CD (compare in **Table 2**).

**Figure 5.**
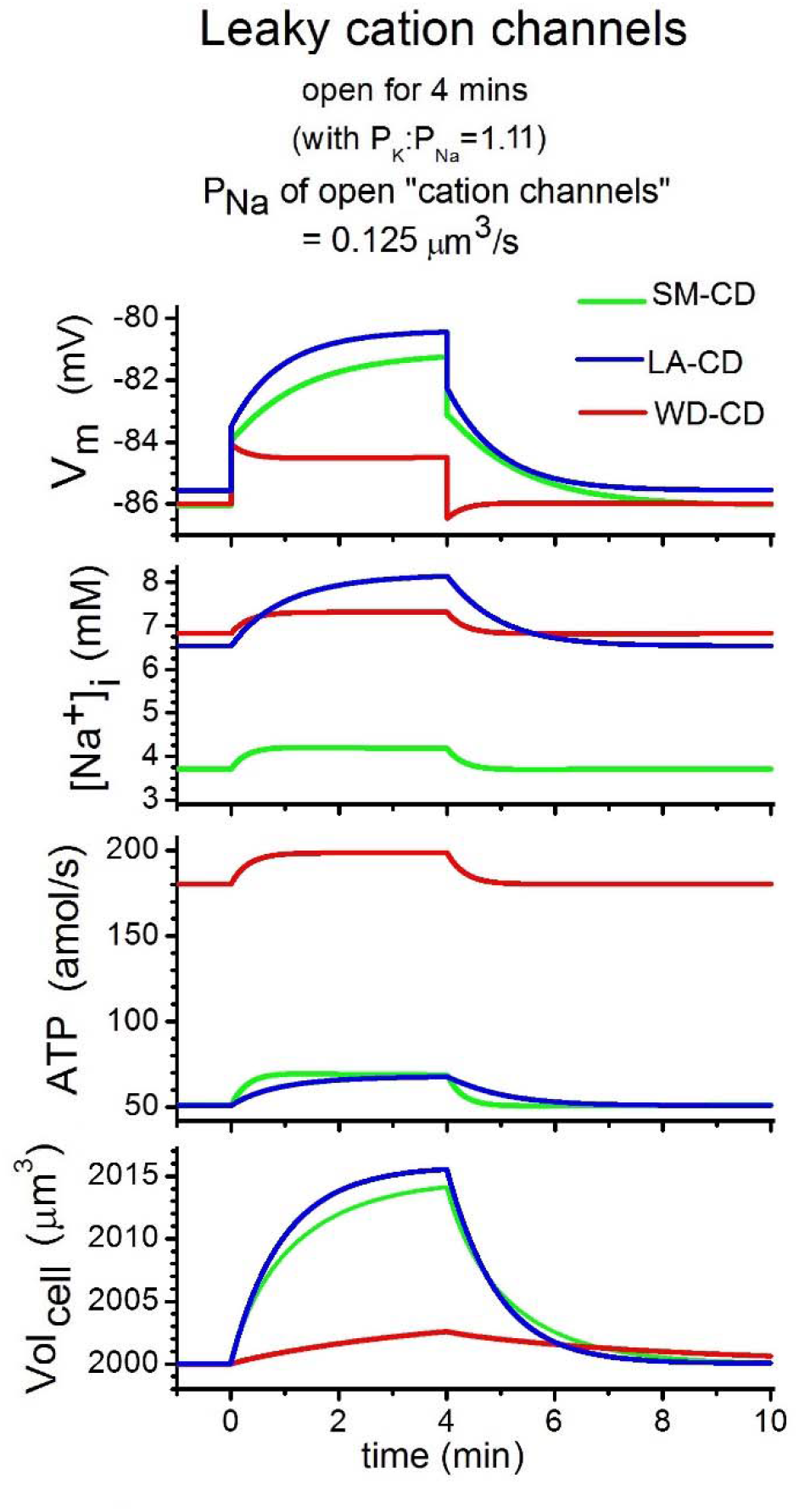
Leaky cation channels. For three minimal P-L/D models (Donnan dominated SM-CD and LA-CD, and Pump-Leak dominated WD-CD), the P-L/D processes are plotted during an identical 4 min step-activation (then step-deactivation) of cation channels (see **Equation 4**). (The on/off would apply if, as suggested by (Lansman 2015), these were mechanosensitive channels rendered hypermechanosensitive (Vandorpe et al 1994) in DMD-damaged sarcolemma, experiencing transient localized membrane stretch. If, more simply, damage induced an open probability increase, there would be an on-step only.) For SM-CD and LA-CD, this increases “resting” P_Na_ by 42% and for WD-CD, by 12%. The V_m_(t) step-on/step-off jumps are “RC” relaxations For all 3 cases V_rest_ is far from E_Na_, near E_K_, and E_Cl_=V_rest_, The depolarizing/hyperpolarizing jumps therefore mostly reflect the onset/offset of inward I_Na_. Slow V_m_(t) changes reflect the active (see ATP-consumption) and passive ion homeostatic feedback fluxes as each model moves toward a new steady-state. Swelling rates are determined by the smaller contributor to net osmotic influx, [Na^+^+Cl^-^]. Since driving forces on Cl^-^ are very small, Volcell changes lag the [Na^+^]_i_ changes even in [big P_Cl_] SM-CD and LA-CD. In Pump-Leak dominated (“neuron-like”) WD-CD, Vol_cell_ changes very little and very slowly due to [small P_Cl_] plus the small Cl^-^ driving force. Cl^-^ influxes, not shown, scale exactly with Vol_cell_ changes.

At 30% pump-strength, LA-CD offers insight for DMD. In both LA-CD and in SM-CD the depolarizations would be readily measurable *in vitro.* But while SM-CD’s attendant ↑[Na^+^]_i_ (~0.5 mM) would not measurably alter “Na^+^-overload”, the rise in LA-CD (to [Na^+^]_i_>8 mM) puts it the range of *mdx* Na^+^-overload (see **Table 1**; item **8**). Note that ~2/3 of the LA-CD Na^+^-overload is attributable, however, to “too little Na^+^-pumping” with ~1/3 from “too much Na^+^-leak”. Increasing cation-leak in SM-CD to 0.250 μm^3^/s (SM-CD-P_Na_ ↑83%) is strongly depolarizing (–86 mV→-77.5 mV), but [Na^+^]_i_ still rises only 3.7→4.5 mM (not shown).

Such calculations indicate that otherwise-healthy fibers (SM-CD) harboring leaky cation channels that doubled P_Na_, would depolarize by ~10 mV, with little ↑[Na^+^]_i_ (<2 mM) or swelling (<1%) whereas DMD-like fibers (LA-CD), though minimally swollen, would become measurably Na^+^-overloaded, depolarized and vulnerable to myogenic discharge.

Another menace from “leaky cation channels” relates to anoxic bouts: the ↑P_Na_ accelerates rundown then compromises recovery (↑I_Naleak_→I_pump_-reserve). In *mdx* fibers stretch-injury can cause prolonged (non-lethal) depolarization (Baumann et al 2020) but the identity of Na^+^-permeant paths rendered leaky by injurious stretch are unknown.

### Chronic ischemia + repetitive stimulation + bleb-damage Na^+^-leaks

Healthy rat soleus fibers can fire 1200 APs (120 Hz, 10 seconds) then briskly return to steadystate (see Clausen, 2015); 100% pump-strength SM-CD recapitulates this stress-test (Joos & Morris submitted) and **Figure 6A** shows that even deeply ischemic LA-CD handle this challenge well, though its recovery is slower by several minutes. The robustness of LA-CD’s response suggests that, *in situ,* comparably ischemic SMFs would have adequate pump-reserve to support Na^+^-transporters since endplate triggered APs would occur at much lower frequencies than the stress-test.

**Figure 6.**
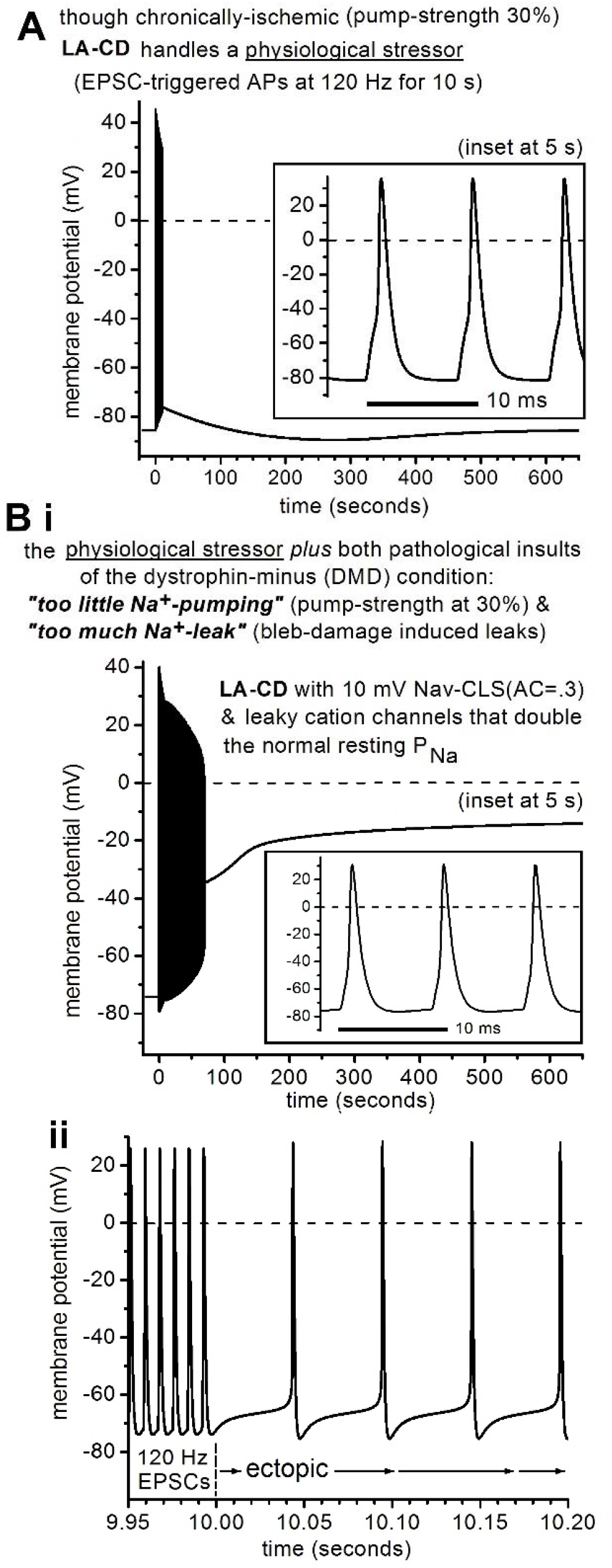
Stress-testing the ischemic model, LA-CD, without/with leaky channels. **A,** V_m_ for LA-CD before/during/after the described stress-test. **Bi**, V_m_ for LA-CD with “bled-damage” (and the resulting Na^+^-leaks as described) us LA-CD before/during/after the same stress-test. **ii**, as the stress-test ends at 10 s, expanded vies of the transition from stimulated to ectopic (“myogenic”) APs.

But dystrophic ischemic fibers might also have bleb-damage Na^+^-leaks. Inflicted with, say, 10 mV Nav-CLS(0.3), or with sufficient cation channel leak to double its P_Na_, LA-CD still passes the stress-test and slowly recovers to its leaky ischemic steady-state (not shown). Robustness, however, has limits as seen in **Figure 6Bi.** There, **i**schemic LA-CD is inflicted with both the above-mentioned Na^+^-leaks then stress-tested. Due to ↑P_Na_, the steady-state (the quiescent state prior to “t=0”) is depolarized by ~12 mV, but the ischemic doubly-traumatized system successfully fires 1200 APs. However, once EPSCs stop at 10 s, (**Figure 6Bii**) it becomes evident that some threshold has been crossed. The system is not on a recovery trajectory. It fires ectopic APs (~20 Hz) of diminishing amplitude; they cease abruptly at ~60 seconds. Overstimulated, ischemic, bleb-damaged LA-CD (still consuming ATP) converges toward a pathological steady-state. P-L/D parameter trajectories (not shown, but as per **Figure 2Ci**) reveal a cytotoxically-swelling system. An excitable-P-L/D bistability scenario has occurred. At some point, while firing 1200 APs, the impaired system destabilized, meaning that it encountered, in parameter space, an unstable threshold beyond which the hyperpolarizing I_pump_ (associated with 30% maximum-strength) was insufficient to counteract total depolarizing I_Naleaks_. *In situ* contracture-induced Ca^2+^-necrosis would likely occur before cytotoxic swelling developed. Slow inactivation of Nav1.4 could help forestall destabilization.

### Na^+^-overload and DMD multiple-jeopardy

Ischemia and bleb-damage typically co-exist for DMD fibers. With ^23^Na-MRI monitoring of patients in mind, we survey (**Figure 7**) their predicted simultaneous impacts on chronic Na^+^-overload. **Figure 7A** (left Y-axis, magenta) plots the smallest pump-strength that ensures SM-CD viability (conservatively defined as “no myogenic firing”) with increasing bleb-damage intensity (X-axis; Nav-CLS(0.3)). The right Y-axis and the inset show the corresponding steady-state [Na^+^]_I_ and Vol_cell_.

**Figure 7.**
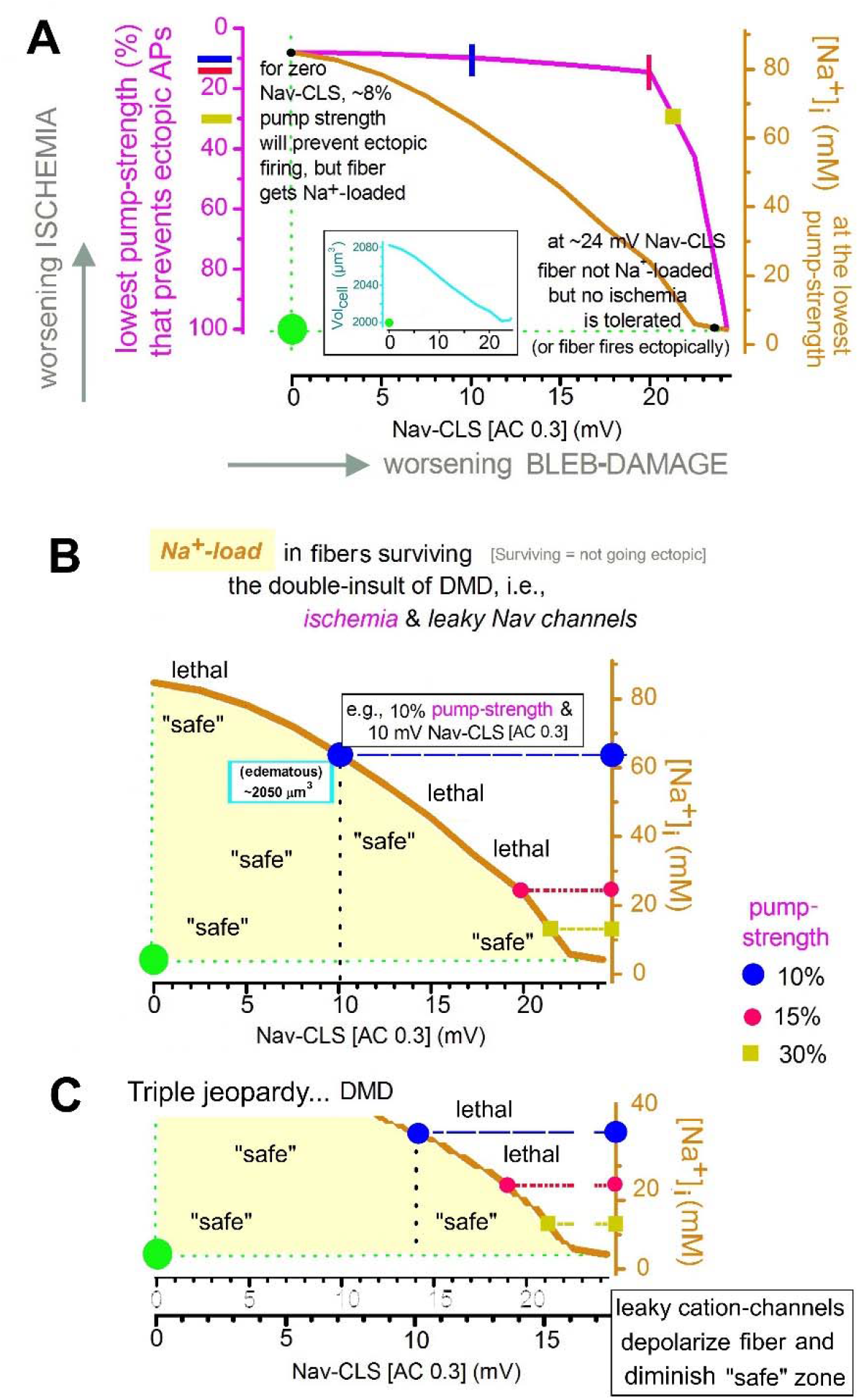
The double insult of DMD: what to expect from ^23^Na-MRI. Myoplasmic Na^+^ levels measured in DMD patients’ muscles by ^23^Na-MRI would represent an average from fibers whose bleb-damage and ischemic severity covered a range (Weber et a 2011, 2012; Gerhalter et al 2019). Likewise for otherwise healthy individuals with muscle injury, e.g., athletes (Hotfiel et al 2018; Dahlmann et al 2016). **(A)** The green dot is the healthy steadystate (no ischemia, no damage, and thus no Na^+^-loading, edema or depolarization). For worsening Nav-CLS (AC=0.3) damage, steady-state [Na^+^]_i_ (and inset, the corresponding steady-state Volcell) at the lowest pump-strength that is able to prevent ectopic firing. **(B)** For the double insult situation, this plot emphasizes that if Na-CLS blebdamage intensity is small, “safe” (versus “lethal”) [Na^+^]_i_-loading is tolerated. **C** is a “sketched” plot as labeled and explained in the text.

**Figure 8.**
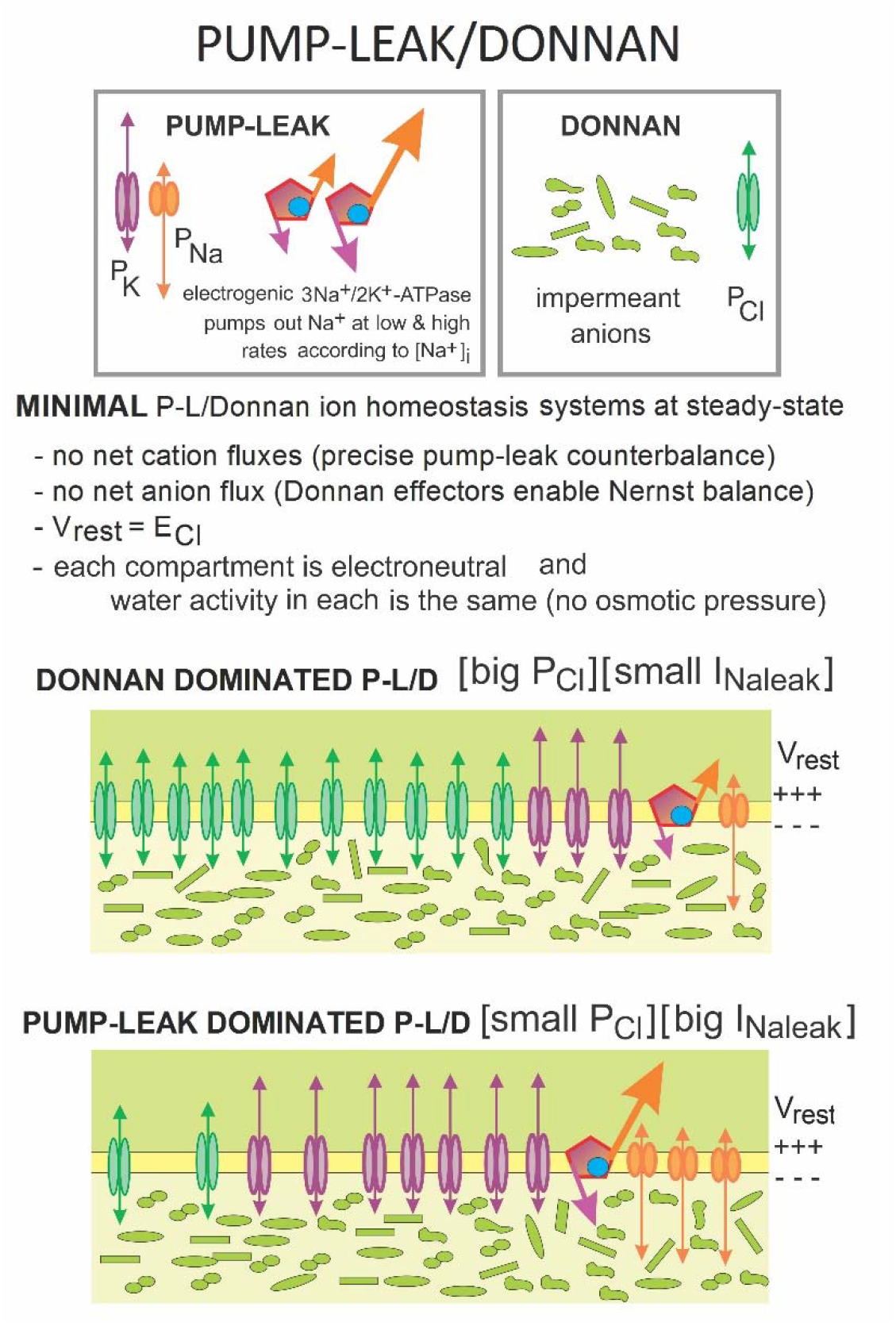
Pump-Leak/Donnan ion homeostasis: two ends of the spectrum. Cytoplasm’s impermeant anions (mostly proteins) are Donnan effectors whose potentially lethal cell-swelling forces are integral to animal cell ion homeostasis, which is a Pump-Leak/Donnan (P-L/D) process. P-L/D processes autonomously maintain and restore (post-ionic perturbations) steady-state [ion]_i(ntracellular_), cell volume (Vol_cell_) and membrane potential (V_m_). In a minimal P-L/D system, steady-state V_m_(=V_rest_) depends on P_Na_:P_K_ and the electrogenicity and [Na^+^]_i_-sensitivity of the pump, 3Na^+^/2K^+^-ATPase. The membrane is H_2_O-permeable and, via P_Cl_, permeable to small anions (here, just Cl^-^). Steady-state Vol_cell_ depends on the molar quantity (not the concentration) of Donnan effectors. SM-CD (the model used here for to represent the P-L/D processes of SMFs) is a minimal P-L/D system to which excitability-machinery is added. During restoration to steady-state following any perturbation, the autonomous P-L/D process can be summarized as follows: 1. the ATP-consuming electrogenic pump “listens” and responds (only) to [Na^+^]_i_ while 2. Passive ion fluxes via P_Cl_, P_Na_, P_K_ “feel” and respond to their electrodiffusive driving forces while standard thermodynamics ensures that no inside/outside H_2_O-activity differential develops. Thus, cytoplasmic electroneutrality and transmembrane osmo-balance are maintained even as the pump (3Na^+^-out/2K^+^-in; net effect, a hyperpolarizing Na) works, rebuilding cellular ion-batteries. Thus, at steady-state in SM-CD, the small hyperpolarizing I_Napump_=the small depolarizing I_Naleak_. These cartoons depict the minimum elements needed for P-L/D steady-states dominated by anion fluxes or cation fluxes, as previously (Joos & Morris, submitted) explained. Whereas SM-CD is a minimal Donnan dominated P-L/D system, CN-CD is a ***non***-minimal Pump-Leak dominated P-L/D system (a K/Cl co-transporter puts its E_Cl_ negative to V_rest_; see footnote, **Table 2**). The Nav and Kv channels (whose m^3^h and n values at V_rest_ are negligible) and the ACh-activated cation channels (AChRs) underlying excitatory synaptic current-trigged APs do not contribute to steady-state, so are not depicted.

**Figure 9.**
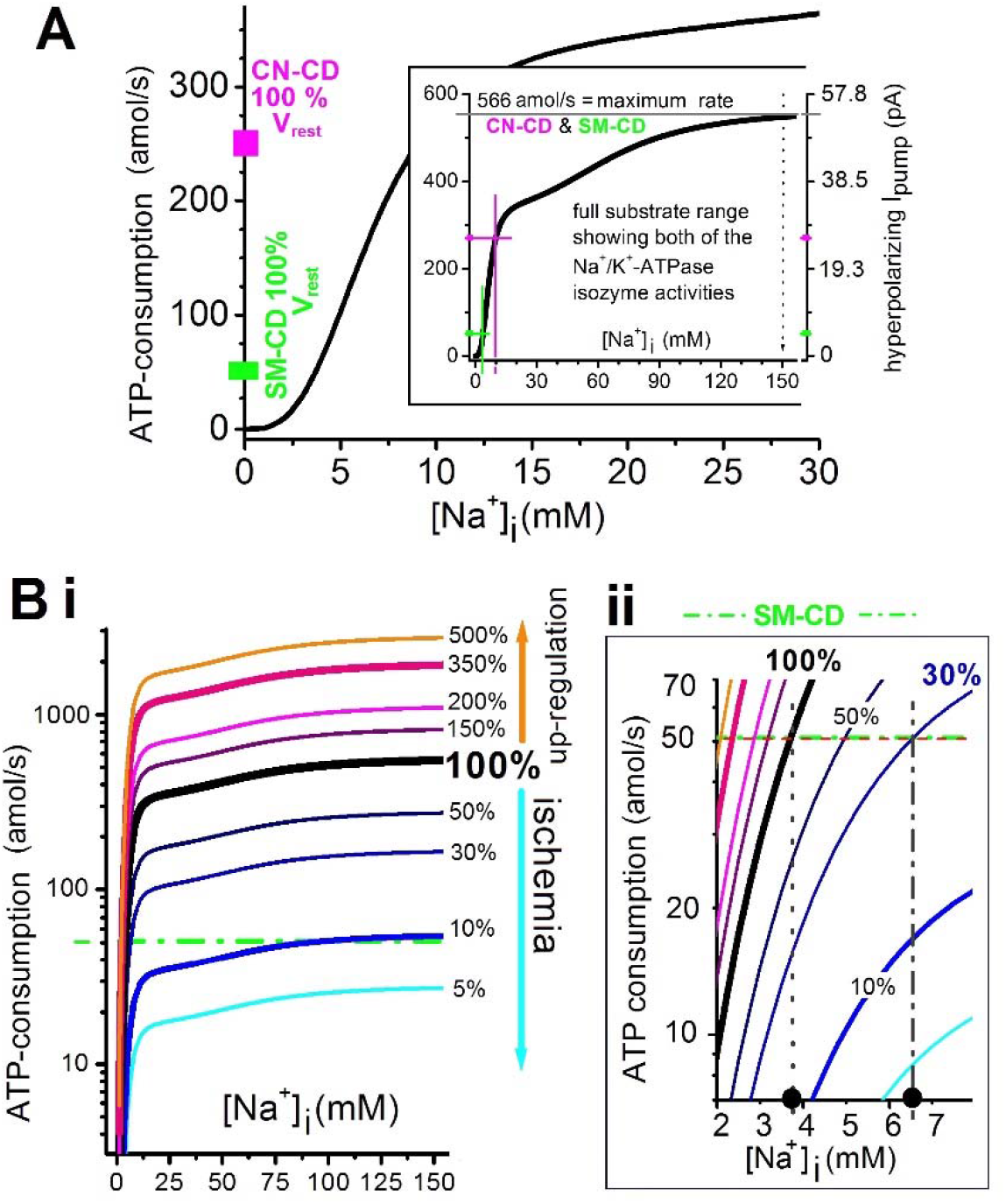
ATP-consumption and [Na^+^_i_. **A.** For all the CD models here, a standalone biochemical descriptor of the pump model (Hamada et al, 2003) i.e., a 3Na^+^/2K^+^-ATPase kinetics plot (substrate: Na^+^i) (see **Equation 10**). The CD models assume a fixed external milieu, so the pump is insensitive to [K^+^]e. Steady-state values for SM-CD and CN-CD are indicated. The right axis shows the equivalent hyperpolarizing electrogenic Ipump (maximally, 54.5 pA). **Bi.** The standard (100%) curve is multiplied by scaling factors to depict ischemia (pump-strengths <100%) and up-regulation (pump-strengths >100%) with logarithmic Y-axes. The dot-dash green line is SM-CD’s resting ATP-consumption. **ii.** expanded lower left sector to show that, notwithstanding almost-identical steady-state ATP-consumption, SM-CD (100% pump-strength) and LA-CD (30% pump-strength) achieve very different [Na^+^]_i_ thanks to steep kinetics in the substrate’s normal physiological range.

**Figure 7B** replots **7A** minus the magenta “pump-strength” curve ([Na^+^]_i_ at 10%, 15%, 30% from **7A** are flagged). The “safe”/“lethal” [Na^+^]_i_ partition-line emphasizes, qualitatively, how bleb-damage reduces a fiber’s tolerance of [Na^+^]_i_-overload. For fibers in DMD-patient muscles, co-ordinates would scatter within the safe zone. Toward the left, a safe fiber near the Na^+^-line would be particularly vulnerable to, say, Ca^2+^-necrosis via reverse Na^+^/Ca^2+^-exchange as per Burr et al (2014). Toward the right, safe fibers would risk excitation-triggered release of Ca^2+^-stores (e.g., Claflin and Brooks, 2008). **Figure 7C** is a “sketch” (no specific computations) depicting how leaky-cation channels further imperil survivability.

Human myoplasmic [Na^+^]_i_ (based on ^23^Na-MRI) is evidently higher than in mice (**Table 1**). Hammon et al (2015) emphasize how ^23^Na-MRI could “be used to gain new insights into {human} Na^+^ homeostasis, presumably leading to better comprehension of pathophysiology”. An anecdotal proof-of-principle study (Dahlmann et al 2016) encourages that view: in a young athlete, muscle injury and recovery (non-injured contralateral=control) was monitored via ^23^Na-MRI. The outcome (in mM) for [Na^+^]_i-control_/[Na^+^]_i_-injured: time=0 18/44; 2 weeks 19/38; 2 months 19/22. Congruent with SM-CD and LA-CD anoxic rundown/recovery (**Figure 2B,Cii,ii,E**), Hammon et al (2015) showed that during anaerobic exercise, muscle exhibits a water-independent (i.e. non-osmotic) ↑[Na^+^]_i_ and, during recovery, a I[Na^+^]_i_. Advances in cell physiological-level instrumentation, too, are will deepen P-L/D investigations of SMFs: a four-electrode method (Heiny et al 2019) now makes it possible to follow multi-ion dynamics and transport processes under voltage-or current-clamp, even monitoring [Cl^-^]_i_, accurately (DiFranco et al 2019). With a flood of observations relevant to DMD fiber ion homeostasis expected, the P-L/D theoretical framework presented here should help make those findings, plus results from the plethora of DMD molecular therapies trails now underway (Verhaart and Aartsma-Rus 2019), more comprehensible, while also suggesting additional lines of inquiry.

## DISCUSSION

Progressive fiber-loss renders DMD patients non-ambulatory by their early teens. 10-15 years later, weakened diaphragm and thoracic muscles cause respiratory failure and premature death (Allen et al 2016; Gerhalter et al 2019). Given the multitude of primary and secondary defects besetting DMD SMFs (e.g.,Murphy et al 2019), fiber survival for weeks let alone decades is remarkable and begs explanation. We therefore mostly ask not “what is wrong with DMD fibers?” but “what is right with DMD fibers?”

An extant model for excitable SMF ion homeostasis (Fraser et al 2011), though powerful and rigorous, is not readily implemented and has seen little uptake for either healthy or ailing fibers. However, guided by the Fraser and Huang (2004) charge difference approach (and SMF permeability ratios) we modified the more recent ion homeostasis model, CN-CD (devised to probe cortical spreading depression (Dijkstra et al 2016)) to obtain SM-CD, our minimal model for ion homeostasis in excitable SMFs (Joos and Morris, submitted). Though both models exhibit patho-physiologically critical (lethal) threshold phenomena (i.e., pump-strength points below which homeostatic insufficiency elicits lethal spontaneous firing), the distinctive Donnan dominated P-L/D strategy of SMFs, safeguards SMFs through short term emergencies and the chronic state-of-emergency that is DMD.

### Overview: SMFs’ Donnan dominated P-L/D process

SMFs’ energy-efficient hyperpolarized steady-state is achieved by a P-L/D process that is Donnan dominated ([big P_Cl_][small I_Naleak_]). To address intermittent overexertion/trauma/misadventure, SMFs maintain an ample pump-reserve. To keep ever-present Donnan swelling forces at bay, SMFs keep P_Na_ extremely small. The resulting [small I_Naleak_] minimizes 3Na^+^/2K^+^-pump expenditures, fosters the ample pump-reserve, keeps anoxic rundown slow and enables the hyperpolarized V_rest_. Given [small I_Naleak_], SMFs can, for no-added-cost, stabilize V_rest_ at or near E_Cl_ by exploiting the low-impedance “battery” available through the [big P_Cl_]/Donnan-effector pairing.

### SMF ion homeostasis ([big P_Cl_[small I_Naleakl_]) confronts DMD as follows

1. Chronic ischemia is tolerated because [small I_Naleak_]→low steady-state ATP-consumption.
2. As ischemic low duty-cycle excitable cells (i.e., mostly at V_rest_), DMD fibers benefit from the no-cost low-impedance [big P_Cl_]/Donnan-effector “battery” that stabilizes their (cheaply) hyperpolarized V_rest_.
3. Bouts of functional ischemia and/or other transient episodes of anoxia/ischemia are survivable because [small I_Naleak_]→(slow rundown) with minimal swelling. How? During rundown, since P_K_>>P_Na_, and V_m_ closely tracks E_Cl_, electroneutrality and osmo-balance are maintained while [Na^+^+Cl^-^+H_2_O]entry stays inconsequential.
4. The chronic Na^+^-overload of DMD is “non-osmotic” in nature (same explanation as in 3).
5. The SMF pump-reserve is so big that even in deeply ischemic (DMD-like 30% pump-strength) SM-CD it exceeds that of 100% “CN-CD neurons” it can support fiber recovery (albeit, more slowly, and within limits) after overstimulation and transient bouts of anoxia and when sarcolemmal damage induces Na^+^-leaks.
6. The large pump-reserve can redress transient the Na-loading associated with the intermittent myogenic firing (triggered, e.g., by sporadic Na^+^-leaks in DMD-damaged sarcolemma).
7. Though pump-reserve “runs out” only at an exceeding low pump-strength (~8% as modeled here), diaphragm fibers (they must regularly fire APs) would destabilize at a somewhat higher pump-strength.
8. “Leaky” Na^+^-permeant channels and ENa-depleting Na^+^-transporters (the latter not modeled here) would exacerbate chronic Na^+^-overload but would not be its sole cause.
9. “Leaky” Nav and cation channels are expected in the bleb-damaged sarcolemma of DMD fibers. Modeling them in the ischemic P-L/D setting shows how they could influence various DMD-patho-phenomena (chronic and/or sporadic depolarization, Na^+^-overload, erratic myogenic firing).
10. Though the Ca^2+^-necrosis conditions of DMD fibers are difficult to pinpoint, they emerge in our SM-CD based analysis as P-L/D thresholds. They are encountered when severe ischemia and/or severe Na^+^-loading (Na^+^-leaks, excess APs) push the P-L/D system to an unstable point in parameter space (i.e., where system-values make it impossible for hyperpolarizing I_pump_ to re-establish a stable physiological state). The emphasis here on “severe” bespeaks the robustness of even deeply compromised DMD fibers. Diverse “only-just” lethally-destabilizing circumstances (see {} brackets) were simulated here, including:

i. (<u>extreme ischemia</u>): 7% pump-strength *{Safe at 9%}.*
ii. (<u>extreme bleb-damage</u>): 100% pump-strength but Nav-CLS(0.3)=30 mV *{Safe at 24 mV}.*
iii. (<u>deep ischemia + over-stimulation + ischemia + membrane damage</u>): 30% pump-strength + APs @120 Hz for 10 s + Nav-CLS(0.3)=30 mV + 2XPNa *{Safe with 1XPNa OR with Nav-CLS(0.3)=0 mV}.*

### An aside

Since a P_Cl_ (not P_K_) dominated resting conductance is mandatory for mammalian SMF resilience, and since some aspects of exertion-related DMD pathology are well-modeled, biologically, via *C elegans* body wall musculature (Hughes et al 2019), we note that resting conductance in nematode muscle is P_Cl_-dominated (Brading & Caldwell, 1971) and that nematodes have ClC channels (Jentsh and Pusch, 2018).

### DMD fibers’ chronic state of emergency

Vertebrate SMFs evolved as long-lived syncytial fibers reliant on the superlative emergency preparedness afforded by a cheaply-maintained pump-reserve far in excess of requirements for normal operation. SMFs thus survive the intermittent ion homeostatic emergencies to which working muscles are prey. These emergencies could involve temporary (minutes to hours) blood-flow restrictions, starvation, over-exertion, tissue trauma, sarcolemmal-damage, sarcolemmal-tearing. Tourniquets are tolerated for prolonged periods (Zhang et al 2020) but toleration of ischemia has a limit. Compartment syndrome (a massive, generally sudden, situation of pressure-impeded blood flow, often compounded by tissue damage) represents an emergency circumstance too extreme even for SMFs, barring surgical intervention that restores blood flow before an irreversible (poorly characterized) threshold is reached (Johnstone and Ball 2019).

DMD fibers, in effect, exist in a state of “chronic emergency”; this is debilitating – exertion is deleterious for DMD patients - yet, thanks to [big P_Cl_][small I_Naleak_] ion homeostasis, non-lethal.

For DMD fibers, I_pumpmax_ is chronically too small and sarcolemmal damage can make operational I_Naleak_ too large. Modeling here repeatedly highlights the need to identity skeletal muscle’s overlooked P_Na_, then to learn if/how it can be safely reduced to counteract these pathological deficits.

The activity regimes of, say, diaphragm versus limb fast-twitch fibers differ markedly. SM-CD is generic. Its clinical utility could be amplified, as appropriate, by modifying pump, leak and excitability characteristics for particular fiber-subtypes. As more molecular, genetic and pharmacologic therapies for DMD come to the clinic (Verart and Aartsma-Rus 2019; Meng et al 2020), non-invasive MRI-monitoring of myoplasmic [Na^+^] and other ion homeostatic indicators (Gerthalter et al 2019) will increasing be used to gauge treatment efficacy. Even in its present utterly basic form, the SM-CD theoretical framework provides fresh insights into the cell physiological meaning of clinical indicators, and helps explain DMD fiber resilience and vulnerability. P-L/D modeling of SMFs facilitates clinical interpretation and generates questions, the most pressing raised here being:

1. What proteins underlie SMFs’ critical small P_Na_?
2. Could DMD fiber longevity be extended by diminishing P_Na_?

## METHODS

### A Pump-Leak/Donnan (P-L/D) model for SMFs

Skeletal muscle fiber (SMF) ion homeostasis is modeled (SM-CD) by a single compartment limited by a semipermeable membrane in an “infinite” (fixed concentrations) extracellular volume (**Table 2**). SM-CD constitutes a single P-L/D “ion homeostatic unit”; an actual multinucleate SMF would be comprised of hundreds or even thousands of such units (depending on sarcolemma area) and, as per Joos and Morris (submitted) would likely have a smaller SA/V (surface area to volume) ratio than SM-CD (thus, slower Δ[ion]_i_ dynamics). The SM-CD membrane encloses a fixed quantity of Donnan effectors: A^-^, impermeant monovalent anions. The extracellular medium has a fixed A^-^ concentration. The membrane is permeable to Na^+^, K^+^, and Cl^-^, ions whose extracellular concentrations are fixed. SM-CD has the same physical characteristics as the Dijkstra et al (2016) neuronal cell, CN-CD (see **Table 2**): a resting cell volume (Vol_cell_) of 2000 μm^3^ and a constant Cm corresponding to a SA of 2000 μm^2^. As such, resting state cells are flaccid (i.e., non-spherical). Lipid bilayers tolerate little lateral expansion, but unless a model-cell inflated to spherical its membrane would not be subject to tension and rupture (see **Table 2**). The permeation pathways of SM-CD and CN-CD (given below) include “resting leak conductances” (i.e., permeation pathways) for 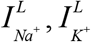, and 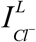 and voltage-gated conductances (permeation pathways) for a transient sodium current, 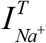, for delayed rectifier potassium current, 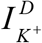, and (CN-CD only) a voltagedependent chloride current, 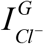. Driving forces acting on ions are, in all cases, electrodiffusive, as depicted by Goldman Hodgkin Katz (GHK) current equations (Hille, 2001). The same 3Na^+^_out_/2K^+^in ATPase pump model (Hamada et al 2003) is used throughout. It produces hyperpolarizing current

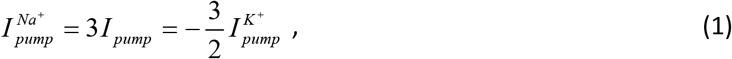

in response to the intracellular [Na^+^] (as per **Figure 4**). In CN-CD only there is a K^+^/Cl^-^ co-transporter for ion flow, *J_KCl_*.

Since animal cells cannot sustain osmotic pressures, intra/extracellular osmolyte concentration inequalities produce a H_2_O flow till osmotic balance is restored; this results in Vol_cell_ changes at rates set by the smaller of the two components giving rise, at any time, to osmotic loading, i.e. to net [Na^+^ + Cl^-^] entry.

### Choice of leak permeabilities

Whereas CN-CD is precisely the Dijkstra et al (2016) model, the SMF version (SM-CD) has a leak permeability ratio 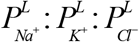 broadly consistent with Fraser and Huang (2004); for more detail see **Setting V_rest_ in SM-CD and other CD models** below.

### GHK driving forces

Currents through open channels (permeability pathways), ion-specific or not, are modeled with the GHK formulation (Hille, 2001). For ion X = Na^+^, K^+^, or Cl^-^, the GHK current is given by:

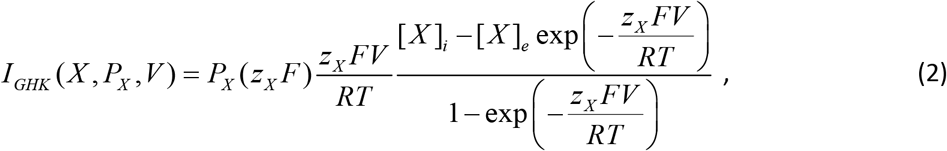

where *P_X_* is the permeability, *z_x_* the valence, and [X]_i_ and [*X*]_e_ the intra- and extra-cellular concentrations of *X* respectively.

### Leak currents

“Leak” permeability mechanisms use the GHK formulation:

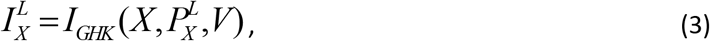

where *X* is either *Na^+^, K^+^* or *Cl^-^* Leak permeability 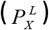 values were adjusted for model variants used here (**Table 2**) in the context of appropriate setting of *V_rest_.*

### Cation channel currents

For non-selective cation channels, we use a P_K_:P_Na_ ratio of 1:1.11 and the formulation:

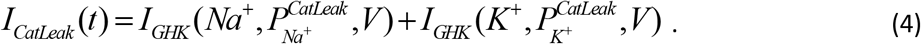

In the present models, cation channel leaks do not contribute to healthy steady-states – they are either transient stimulatory currents through SMF-endplate type AChR channels, or pathological leaks (hence “leaky” cation channels).

### Transient voltage-gated Na^+^ current

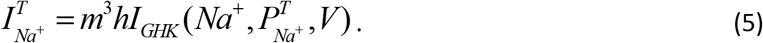

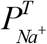 is the maximal membrane permeability to Na^+^ through a V-gated channel (operating in a Hodgkin-Huxley (H-H) fashion). *m* is the H-H Na^+^ channel activation/deactivation gating variable and *h* is the H-H Na^+^ channel inactivation/recovery gating variable. The current’s driving force also follows the GHK form of **Equation 1**.

### Delayed rectifier K^+^ current

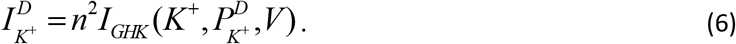

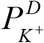 is the maximal membrane permeability of K^+^ through a V-gated channel (operating in a H-H fashion); *n* is the delayed rectifier K^+^ channel activation/deactivation gate variable.

### Voltage-dependent gating

The non-dimensional gating parameters *m, h, n* evolve in time according to:

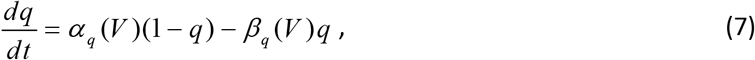

where *q* ∈ {*m, h, n*} are as defined above, and the voltage dependent α_q_(*V*) and β_q_(*V*) refer, for *m and n* to gate activation and deactivation and, for *h* to inactivation and recovery from inactivation. **Table 3** gives the voltage dependences for the relevant rate constants.

**TABLE 3.** Voltage-dependent rates for H-H gating variables*

| Term | Expression | Description |
| --- | --- | --- |
| $\alpha_m(V)$ | $\frac{0.32(V + 52 \text{ mV})}{1 - \exp\left(-\frac{V + 52 \text{ mV}}{4 \text{ mV}}\right)} \frac{\text{kHz}}{\text{mV}}$ | Activation rate for transient Na<sup>+</sup> channel |
| $\beta_m(V)$ | $\frac{0.28(V + 25 \text{ mV})}{\exp\left(-\frac{V + 25 \text{ mV}}{5 \text{ mV}}\right) - 1} \frac{\text{kHz}}{\text{mV}}$ | Deactivation rate for transient Na<sup>+</sup> channel |
| $\alpha_h(V)$ | $0.128 \exp\left(-\frac{V + 53 \text{ mV}}{18 \text{ mV}}\right) \text{ kHz}$ | Inactivation rate for transient Na<sup>+</sup> channel |
| $\beta_h(V)$ | $\frac{4}{1 + \exp\left(-\frac{V + 30 \text{ mV}}{5 \text{ mV}}\right)} \text{ kHz}$ | Recovery from inactivation rate for transient Na<sup>+</sup> channel |
| $\alpha_n(V)$ | $\frac{0.016(V + 35 \text{ mV})}{1 - \exp\left(-\frac{V + 35 \text{ mV}}{5 \text{ mV}}\right)} \frac{\text{kHz}}{\text{mV}}$ | Activation rate for delayed rectifier K<sup>+</sup> channel |
| $\beta_n(V)$ | $0.25 \exp\left(-\frac{V + 50 \text{ mV}}{40 \text{ mV}}\right) \text{ kHz}$ | Deactivation rate for delayed rectifier K<sup>+</sup> channel |

\* Based on Kager et al (2000) but as used in Dijkstra et al (2016) with one typographical error corrected.

### Voltage-dependent Cl^-^ current

The SLC26A11 ion exchanger based voltage-dependent Cl^-^ conductance (Rungta et al, 2015) is as described by Dijkstra et al (2016):

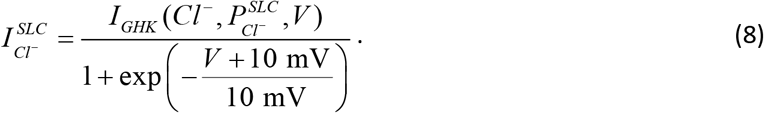

Except in CN-CD and MN-CD, this pathological cortical neuron specific conductance is set at zero.

### K^+^/Cl^-^ cotransporter

This cotransporter (strength *U_KCl_*) is present only in CN-CD (it is not evident in skeletal muscle; Pedersen et al 2016) where (as per Dijkstra et al 2016) it is given by

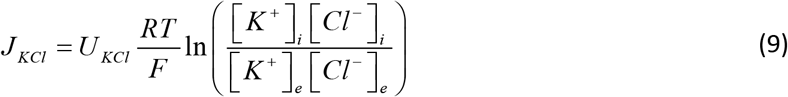

(except that there is an error in the log term of Eq. (6) in Dijkstra et al 2016).

### 3Na^+^/2K^+^ ATPase pump current

The electrogenic 3Na^+^(out)/2K^+^(in) ATPase modeled here is that of Hamada et al (2003):

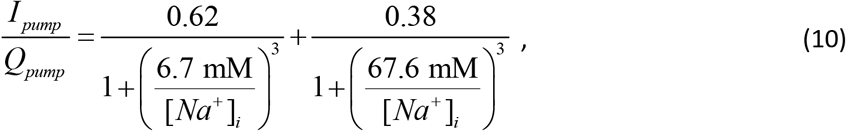

where

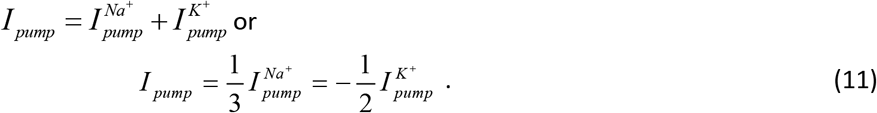

Thus *I_pump_* signifies a net hyperpolarizing Na^+^ outflow. The Hamada et al (2003) formulation is appropriate for models that assume an invariant extracellular medium. It depicts the pump’s two intracellular Na^+^-binding sites (with ~10-fold different affinities) but has no term for the extracellular K^+^*-* binding site. Here, pump-strength (*Q_pump_*) is varied in many computations (e.g., it is set to zero to depict anoxia, diminished towards zero to depict ischemia and multiplied for up-regulation).

### ATP-Consumption

ATP-consumption by the Na^+^/K^+^ ATPase pump in the all the CD models is given as

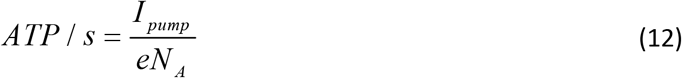

where *e* is the electronic charge (1.6 x 10^-19^Coul) and *N_A_* is the Avogadro number. In other words, *Ipump* of 1 pA is equivalent to an ATP-consumption of 10.38 amol/s.

### Cell Volume

Cellular swelling is driven by an influx of water at rates that depend on the transmembrane osmotic gradient. The rate of change of cell volume, Vol_cell_, due to water influx is given by:

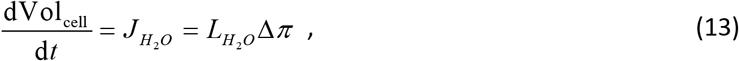

where Δ*π* = *RT* {[*S*]_*i*_. –[*S*]_*e*_), and [*S*]_*i*_ and [*S*]_*e*_ denote total concentrations of intra- and extracellular solutes and *L_H_2_O_* is the effective membrane water permeability. This equilibration is typically assumed to be nearly instantaneous relative to the ion flows (Fraser and Huang, 2004; Dijkstra et al 2016; Kay, 2017; though see also Dmitriev et al 2019). Accordingly, in modeling here, osmotic ion fluxes, not the H_2_O fluxes what limit the rate of osmotic swelling or shrinkage.

### Nernst potentials

The equilibrium potential (Nernst potential) for each ion

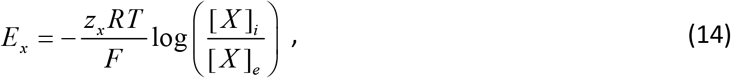

where *X* is either Na^+^, K^+^, or Cl^-^ ions and *Z_X_* is the valence of each ion.

### Charge difference and membrane potential

Because CD models keep track of the absolute number of ions flowing across the cell membrane (of capacitance *C*_m_) no differential equation is needed for *V*_m_. Instead, an accounting equation is used, made simple because the extracellular space is kept neutral:

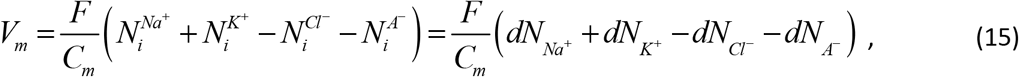

where 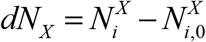 is the difference in the number of ions of species *x*, between its present value 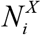 and a reference value 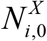. The reference values are those yielding *V_m_* = 0 V, which is equivalent to a neutral intracellular space 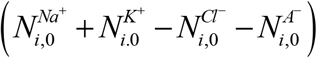. Because *N_A-_* is constant, note that *dN_A-_* = 0 in **Equation 15.** Number differences, *dN_x_*, are easier to work with than total numbers of ions *N_x_* (or than concentrations). For typical cell C_m_ values, voltages in the mV range correspond to net intracellular charge differences (measured as a number of singly charged ions) of the order of attomoles (10^-18^ moles). For instance, when *V*_m_ varies in the range [-100mV, 100mV] this corresponds to a change in net singly charged ions of [-20.7, +20.7] amol. With our *Vol*_cell_ = 2000 μm^3^, this yields tiny concentration changes [-0.01, 0.01] mM. For solutions whose concentrations are in the 1-100 mM range, there could be 4-5 orders of magnitude difference between steady-state values and the changes (in the 0.0001 to 0.01 mM range). Computations based on concentrations, therefore, tend to be unstable. For this reason, calculations in this study are based on changes in ion numbers (in amol units).

### Number of intracellular ions

CD models account for the change in intracellular ions at any given moment (Fraser and Huang 2004, 2007; Dijkstra et al 2016), via the following simple relationships of the respective currents:

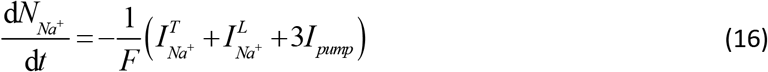

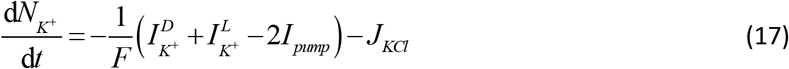

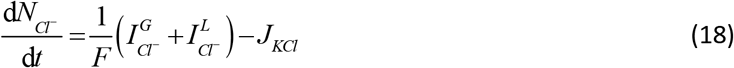

### Setting V_rest_ in SM-CD and other CD models

Experimentally, excitable cells’ *V_rest_* (i.e., steady-state *V_m_*) values are generally more accessible than cytoplasmic ion concentrations or cell volume, making it useful to anchor a model with a consensus *V_rest_* value. For SM-CD, we chose V_rest_= −86, with parameter determinants established in an iterative process as follows: first, a number is chosen for impermeant anions 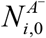, consistent with the system’s total cation concentration (as per the extracellular solution). We started with the CN-CD value, knowing that fine-tuning would be needed to meet our (self-imposed) requirement that SM-CD have the same resting Vol_cell_ as CN-CD. Thus, note in **Table 2** the slightly different 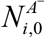 for CN-CD and SM-CD (likewise, LA-CD and SM-CD, to give identical resting Vol_cell_).

At rest, Na^+^ and K^+^ leak currents (for a system with a given pump-strength), must precisely balance. In other words, *V_m_* converges (along with ion concentrations and Vol_cell_) on steady-state (=*V*_rest_) when

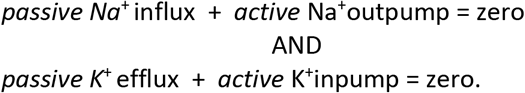

For the 3Na^+^/2K^+^ ATPase pump current, this requirement is met when:

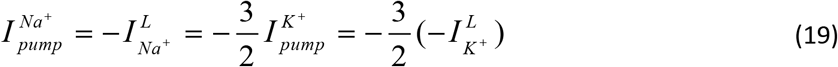

For P-L/D systems with a given pump-strength, *V_rest_* varies monotonically with P_Na_:P_K_ ratio; low ratios yield hyperpolarized *V_rest_* values, high ratios, depolarized ones. *V_rest_*=-86 mV for SM-CD requires P_Na_:P_K_ = 0.03: 1 (likewise for WD-CD). Absolute values for P_Na_ and P_K_, and for P_Cl_ (**Table 2**) were guided by the P_Na_:P_K_:P_Cl_ ratio (0.02: 1: 3) reported for amphibian skeletal muscle (Fraser and Huang 2004). For absolute P values, a straightforward constraint was the choice to give SM-CM the same area (Cm) as CN-CD.

As per **Equation 16**, a pump stoichiometry other than 3Na^+^out/2K^+^in would, all else being equal alter V_rest_. Pump stoichiometry is invariant here, but see Dmitriev et al (2019).

### Excitability and safety factor for SM-CD

For SM-CD to be hyperpolarized and appropriately excitable (i.e. “relatively inexcitable”), its input impedance had to be a) notably less than for CN-CD and b) predominantly P_Cl_-based (Pedersen et al 2016). With resting-P values set, the need to trigger spikes near −60 mV (Fu et al 2011) with a reasonable-sized safety factor (Ruff 2011) had to be met. To achieve safety factor ~1.5, Nav and Kv “densities” in SM-CD were set at 3X the CN-CD level (a larger P_Cl_ would have required even greater V-gated channel densities). Thus, absolute P_Cl_, P_Na_ and P_K_ values of SM-CD are biologically appropriate, but leave room for physiological modulation (to, say, alter V_rest_ via ΔP_Na_ or ΔP_K_, or to modulate excitability by ΔP_Cl_).

**Table 2** shows that m^3^h is vanishingly small at V_rest_ in SM-CD; for CN-CD it adds an extremely small Nav channel contribution to the operational value of P_Na_ that negligibly affects V_rest_.

### Cytoplasmic Donnan effectors

Once V_rest_ is set via P_Na_ and P_K_, the intracellular anion concentrations are determined uniquely for the resting state. If Cl^-^ transport is purely passive (i.e., no involvement of secondary transport) as in SM-CD, then E_Cl_ = V_rest_ and therefore:

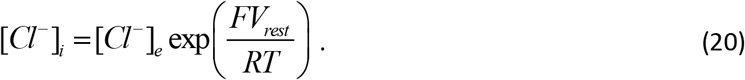

[A^-^]_i_ then follows from the voltage and osmotic balance requirements.

The voltage requirement yields:

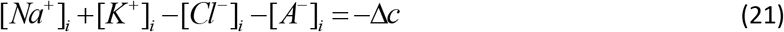

where Δc is the tiny excess concentration of anions associated with V_re_st and equal to [intracellular anions]-[intracellular cations]:

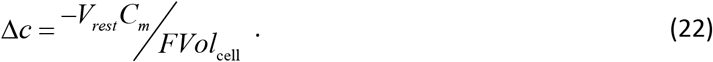

Δc = 0.00886 mM for *V_rest_* = −86 mV. And the osmotic equilibrium condition is:

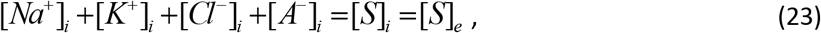

where [S]_I_ and [S]_e_ are defined below **Equation 13**. **Equations 21** and **23** yield:

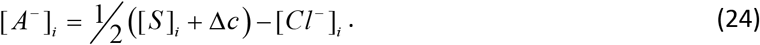

A P-L/D model-cell’s design for steady-state imposes its [A^-^]_i_. Cell volume adjusts to reflect that constraint 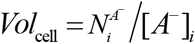. As mentioned above, SM-CD’s 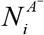 was set to give resting Vol_cell_ =2000 μm^3^, i.e. 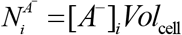. Here, the quantity of impermeant anions 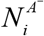 is invariant, but for many *in vivo* circumstances, it would vary, and variations in Vol_cell_ would be expected in such cases.

### Instantaneous perturbations

Experimental solution changes (e.g., as in brain slice experiments) typically require finite “wash-in/wash-out” times. Dijkstra et al (2016) mimicked such solution changes (affecting pump rates and channel gating etc) but here, doing so would have unnecessarily obscured mechanistic underpinnings of responses. Thus, pump-off (anoxia) and pump-on (restoration of pump-strength) changes and channel gating changes (Nav and cation channels open probabilities) are instantaneous.

### Maximum cell volume before lysis

Both CN-CD and SM-CD have C_m_=20pF and steady-state Vol_cell_=2000 μm^3^. If 0.01F/m^2^ (= 0.01 pF/μm^2^) is the specific capacitance of the bilayer, membrane area is 20/0.01 = 2000 μm^2^. With 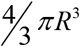 the volume of a spherical cell and *4πR^2^* its surface area, maximum Vol_cell_ as the cell swells (to spherical) would be =4/3π(2000/4π)^3/2^ = 8410.4 μm^3^. Given a 4% bilayer elasticity strain limit (yielding membrane area = 2080 μm^2^), rupture would occur at 8920 μm^3^. Thus, in bifurcation plots, the notional Donnan Equilibrium (DE) values indicated for reference, are unachievable by these models. Note too that present models depict neither surface area regulation nor membrane tension homeostasis (see Morris 2018).

### Excitatory post-synaptic current (EPSC) via AChR channels

SM-CD action potentials are initiated by EPSCs, i.e. macroscopic end-plate currents through (acetylcholine receptors) AChRs, which are non-selective cation channels that pass Na^+^ and K^+^ as per the GHK formalism. The EPSC time course mimics the *g*(*t*) reported in Wang et al (2004). As reported in Hille (2001), 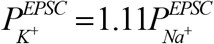. The function *g*(*t*) has a maximum of 1 and 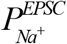 has a 1.5-fold safety factor (i.e., an amplitude adjusted to 1.5X the threshold required to elicit an AP in SM-CD).

The end-plate current is:

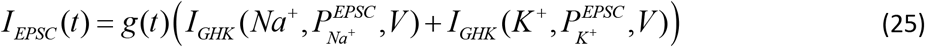

### Nav-CLS depiction of bleb-damage to Nav-bearing membrane

The extent (in mV) of Nav-CLS as depicted in **Figure 1D** increases with the intensity of bleb-damage (Boucher et al 2012; Morris 2012; Morris et al 2012; Joos et al 2018). Cell survival seems improbable with 100% of the Nav-bearing membrane mechanically bleb-damaged. Arbitrarily, therefore, we model damage to 30% of the Nav population [i.e. **A**ffected **C**hannels, AC=0.3]; “damage”, imposed at graded intensities, is “Nav-CLS(0.3) mV”.

The voltage dependences of the rate constants α_q_ (V) and β_q_ (V) for *m* and *h* are the same as in **Equation 7** but shifted by voltage *LS* (in mV), the **L**eft **S**hift. Therefore, for the Affected Nav Channels (AC), *m* and *h* evolve according to:

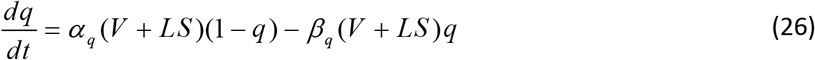

where *q* is either *m* or *h*.

### Computational methods

Calculations involved solving sets of first order differential equations. These were done using Python with the ordinary differential equation solver *odeint.*

## ACKNOWLEDGEMENTS

We acknowledge financial support from Natural Sciences and Engineering Council (Canada) (BJ and JJW) and support from the Ottawa Health Research Institute (CEM).

